# Multiplex imaging of quantal glutamate release and presynaptic Ca^2+^ at multiple synapses *in situ*

**DOI:** 10.1101/336891

**Authors:** Thomas P. Jensen, Kaiyu Zheng, Nicholas Cole, Jonathan S. Marvin, Loren L. Looger, Dmitri A. Rusakov

## Abstract

Information processing by brain circuits depends on Ca^2+^-dependent, stochastic release of the excitatory neurotransmitter glutamate. Optical glutamate sensors have enabled detection of evoked and spontaneous synaptic discharges. However, monitoring presynaptic function and its underpinning machinery *in situ* requires simultaneous readout of quantal glutamate release and nanomolar presynaptic Ca^2+^. Here, we find that the fluorescence lifetime of the red-shifted Ca^2+^ indicator Cal-590 is Ca^2+^-sensitive in the nanomolar range, and employ it in combination with green glutamate sensors to relate quantal neurotransmission to presynaptic Ca^2+^ kinetics. Imaging of multiple synapses in an identified neural circuit reveals that fluctuations both in spike-evoked Ca^2+^ transients and in resting presynaptic Ca^2+^ can affect release efficacy. At the sub-microscopic level within individual presynaptic boutons, we detected no consistent co-localisation of presynaptic Ca^2+^ entry and glutamate release sites, suggesting loose coupling between the two. The present approach broadens qualitatively our horizon in understanding release machinery of central synapses.

---

Stochastic, Ca^2+^-dependent release of the excitatory neurotransmitter glutamate by individual synapses underpins information handling and storage by neural networks. However, in many central circuits glutamate release occurs with a low probability and a high degree of heterogeneity among synapses ^1, 2^. Therefore, methods to probe presynaptic function in an intact brain aim to reliably detect presynaptic action potentials (APs), record the presynaptic Ca^2+^ dynamics, and register release of individual glutamate quanta with high temporal resolution and broad dynamic range. The optical quantal analysis method went some way toward this goal, by providing quantification of release probability at individual synapses in brain slices ^3, 4^. In parallel, advances in the imaging techniques suited to monitor membrane retrieval at presynaptic terminals have enabled detection of synaptic vesicle exocytosis in cultured neurons ^5^. Recently developed optical glutamate sensors ^6, 7^ have drastically expanded the sensitivity and the dynamic range of glutamate discharge detection in organised brain tissue ^8^. However, such methods on their own cannot relate neurotransmitter release to presynaptic Ca^2+^ dynamics, which is the key to understanding presynaptic release machinery. Direct probing of this relationship has hitherto been feasible only in the elegant studies of giant synapses that permit direct electrophysiological access ^9–11^.

Furthermore, conventional methods of optical detection based on fluorescence intensity measures face multiple challenges when applied in turbid media such as organised brain tissue. Registered emission intensity can be strongly affected by focal drift, photobleaching, or any experimental concomitants (such as cell swelling or temperature fluctuations) impacting on light scattering. To overcome such difficulties in Ca^2+^ imaging, we have recently developed a technique to employ Ca^2+^-sensitive fluorescence lifetime imaging (FLIM) of the Ca^2+^ indicator OGB-1 ^8, 12, 13^. The FLIM readout provides nanomolar-range Ca^2+^ sensitivity and is insensitive to light scattering, dye concentration, focus drift or photobleaching. However, OGB-1 emission is chromatically inseparable from that of existing glutamate sensors, thus prohibiting simultaneous imaging of glutamate release ^8^. To deal with this challenge, we have systematically explored time-resolved fluorescence properties of Ca^2+^ indicators and discovered that the fluorescence lifetime of red-shifted Cal-590, an indicator that has been successfully used in deep-brain imaging ^14^, is also sensitive to low nanomolar Ca^2+^.

In parallel, we have developed a glutamate sensor variant SF-iGluSnFR.A184S, which has a suitable dynamic range but a slower off-rate than the existing SF-iGluSnFR.A184V, and thus permits monitoring of released glutamate quanta simultaneously at multiple release sites. We used organotypic brain slice preparations to image chromatically separated concurrent signals reporting evoked glutamate release and presynaptic Ca^2+^ kinetics, at individual or multiple synapses, recorded simultaneously or sequentially, in cell axons traced from the soma. This multiplex imaging enabled us to establish some basic relationships between nanomolar presynaptic resting Ca^2+^, AP-evoked Ca^2+^ entry, and synaptic release properties. At the sub-microscopic level within individual axonal boutons, we took advantage of this method to evaluate localisation of glutamate release sites and their possible co-localisation with presynaptic Ca^2+^ entry, understanding of which is fundamental to neurobiology ^15, 16^. The present approach thus opens a new horizon in monitoring synaptic release function, its use-dependent changes, and the underpinning Ca^2+^ machinery in identified brain circuits *in situ*.

## RESULTS

### FLIM readout of red-shifted Cal-590 reports low presynaptic Ca^2+^

In search of suitable red-shifted Ca^2+^ indicators we tested and excluded the classical rhodamine-based dyes: they appear highly lipophilic, a property undesirable for imaging in axons where free cytosolic diffusion is required to obtain sufficient indicator level away from the soma. Because the fluorescein-based, red shifted Cal-590 was previously found to provide reliable Ca^2+^ monitoring for deep brain imaging *in vivo* ^14^, we set about testing the Ca^2+^ sensitivity of its fluorescence lifetime, particularly for two-photon excitation at λ_x_^2p^ ~910 nm, alongside the green glutamate sensor iGluSnFR ^6, 17^. We discovered a clear dependence between the Cal-590 fluorescence lifetime and [Ca^2+^] using a series of Ca^2+^-clamped solutions, as described previously ^12^ and recently outlined in detail ^13^ (Fig. 1a; Methods). This dependence was most pronounced around λ_x_^2p^ ~ 910 nm (Supplementary Fig. 1a) and appeared largely insensitive to temperature (and associated viscosity changes) between 20-37°C (Supplementary Fig. 1b). Several other tested Ca^2+^ indicators, including red-shifted Asante Calcium Red and Calcium Ruby-nano, showed no usable [Ca^2+^] sensitivity in their fluorescence lifetime (Supplementary Fig. 1c).

Aiming to calibrate Cal-590 lifetime for [Ca^2+^], we used the protocol established previously for the green Ca^2+^ indicator OGB-1 ^12, 13^. The procedure employs the Normalised Total Count (NTC) method in which photon counts are integrated, as the area under the life time decay curve, over the Ca^2+^-sensitive time segment (in this case ~3 ns post-pulse) and the result is related to the peak fluorescence value (Fig. 1a, Methods). This ratiometric method substantially lowers the minimum requirement for the photon counts compared to traditional multi-exponential fitting in FLIM, which in turn shortens minimal acquisition time and improves spatiotemporal resolution of Ca^2+^ imaging ^12, 13^. The calibration outcome best-fitted with the logistic function showed the greatest sensitivity of the Cal-590 FLIM readout in the 0-200 nM range (Fig. 1b).

**Figure 1.**
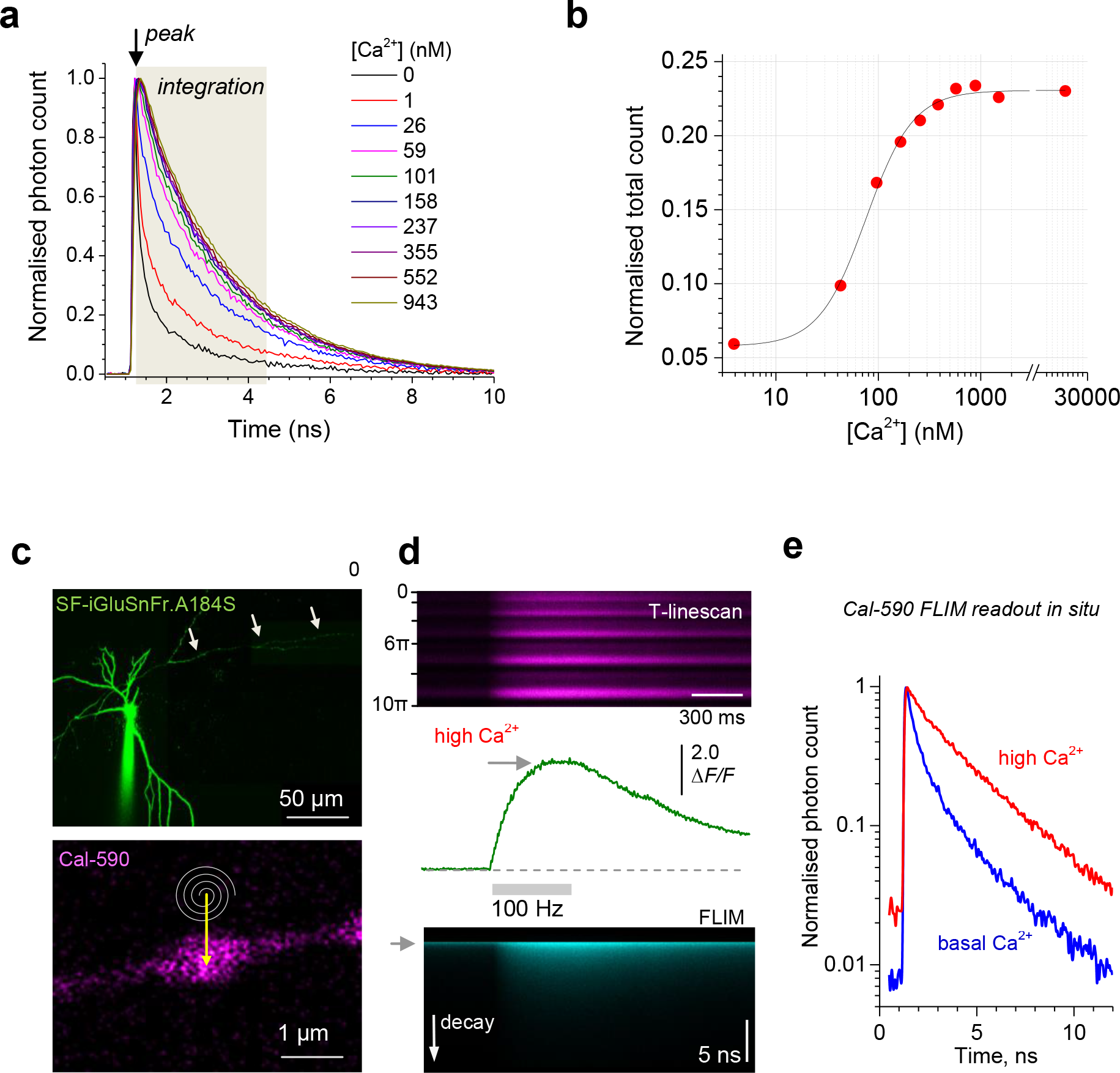
Fluorescence lifetime of Cal-590 provides readout of low intracellular Ca^2+^. **(a)** Fluorescence lifetime decay curves of Cal-590 in a series of calibrated [Ca^2+^]-clamped solutions finely adjusted to include appropriate intracellular ingredients ^12, 13^ (Methods). Fluorescence lifetime traces are normalised to their peak values (at ~1.3 ns post-pulse); the area under the curve over the time interval of 3 ns (tan shade) was measured and related to the peak value thus providing Normalised Total Count (NTC) readout; λ_x_^2p^ = 910 nm; temperature 33°C. **(b)** The Cal-590 FLIM [Ca^2+^] sensitivity estimator, normalised total count, fitted with sigmoid type function (χ^2^ = 2.11·10^-5^, R^2^ = 0.996). **(c)** *Top:* Example of a CA3 pyramidal cell (iGluSnFR channel); arrowheads, proximal part of the traced axon; patch pipette is seen. *Bottom:* Example of an axonal bouton (Cal-590 channel) traced from the cell soma; spiral line and arrow, tornado linescan applied in the middle of the bouton. **(d)** Example of a single-bouton Cal-590 signal during a 500 ms 100 Hz burst of spike-inducing somatic 1 ms current pulses: recorded as a tornado linescan (*top*; ordinate, spiral rotation angle 0-10π reflects five concentric spiral circles, 2π radian/ 360° each), fluorescence intensity (integrated over the 10π spiral scan) time course (*middle*) and fluorescence decay (FLIM, *bottom*; ordinate, decay time; grey arrowhead, laser pulse onset). **(e)** Intra-bouton Cal-590 fluorescence decay time course (normalised to peak) representing basal [Ca^2+^] (blue) and peak [Ca^2+^] (red), in the experiment shown in **d**.

To verify that this approach could be applied in axons traced in organised brain tissue, we turned to organotypic hippocampal brain slices. Because Cal-590 could not be reliably used for axon tracing (due to its weak emission at low baseline [Ca^2+^]) we employed the bright fluorescence of the optical glutamate sensor iGluSnFR: two sensor variants, A184V and A184S, provided similarly bright expression labelling. We achieved sparse cell expression by biolistically transfecting iGluSnFR (Methods), patch-loaded iGluSnFR-expressing CA3 pyramidal cells with Cal-590 (300 μM), and traced their axonal boutons for at least ~200 microns from the cell soma, towards area CA1 (Fig. 1c). The axonal bouton of interest was imaged using a spiral (‘tornado’) scanning mode (Fig. 1c, bottom), which we showed previously to enable rapid (1-2 ms) coverage of the visible bouton area during single sweeps ^8^. Next, we tested whether the dynamic range of Cal-590 FLIM readout inside axons *in situ* was similar to that in calibration conditions (Fig. 1a-b). We therefore recorded both intensity and lifetime decay of the axonal signal, first, in low Ca^2+^ resting conditions (for ~400 ms), and secondly, following a 500 ms 100 Hz burst of spikes evoked at the soma (Fig. 1d), which should reliably saturate the indicator ^18, 19^. Comparing the respective Cal-590 fluorescence decay curves confirmed a wide dynamic range of [Ca^2+^] sensitivity *in situ* (Fig. 1e), consistent with the calibration data *in vitro* (Fig. 1a-b).

### Glutamate sensor SF-iGluSnFR.A184S enables monitoring of quantal glutamate release simultaneously at multiple synapses

We next asked whether the kinetic features of the iGluSnFR sensors were suited for simultaneous imaging of multiple axonal boutons. This consideration was important because the laser beam has to dwell for 1-2 ms at each bouton to achieve signal-to-noise ratio comparable to that obtained under tornado-scanning of individual boutons ^8^. Thus, reliable imaging of fluorescent signals concurrently at 4-5 locations requires the signals to last (preferably near its peak) for at least 10-15 ms. Fortuitously, this requirement is normally met by high-affinity Ca^2+^ indicators including Cal-590 ^14^.

We noted that the recently developed glutamate sensor variant SF-iGluSnFR.A184S ^17^ had a much lower off-rate (slower fluorescence decay) than the faster variant SF-iGluSnFR.A184V (Supplementary Fig. 2a). To test whether the A184S variant was suited to reliably report release of glutamate simultaneously from several axonal boutons, we implemented a scanning mode in which the laser beam travelled, in a cycling manner, across several boutons dwelling at each bouton for 1-1.5 ms (Fig. 2a; Supplementary Fig. 2b). The resulting datasets were re-arranged so that the fluorescence dynamics at individual boutons were represented by ‘pseudo-linescans’ with a time step of 5-7 ms (Fig. 2b; Supplementary Fig. 2c). The A184S indicator provided stable recordings over multiple trials, also revealing no glutamate signal cross-talk between neighbouring boutons (Supplementary Fig. 2d). These settings enabled direct readout of release probability and, at least in some cases, evaluation of the quantal content of release at several synapses simultaneously. Quantal analyses were carried out by adapting a classical approach in which the signal amplitude histogram (SF-iGluSnFR.A184S *ΔF/F* readouts) was systematically fitted with multiple Gaussians, with the fitting parameters constrained by the measured dispersion of the signal noise (release failure response) and by the initial-condition range of peak locations ^20^, upon which it carried on unsupervised (Methods). The method revealed multiple, nearly equally-spaced peaks in the amplitude histogram, thus indicating the numbers and sizes of released neurotransmitter quanta (up to three quanta per release; Fig. 2c-d; Supplementary Fig. 2e-f; the reliability of quantal assessment can be further improved with larger trial numbers). Finally, it was also possible to perform multiplex imaging of presynaptic Ca^2+^ dynamics (using Cal-590 fluorescence) and glutamate release in similar settings with the A184S variant (Supplementary Fig. 3). However, in our hands the relatively slow kinetics of iGluSnFR.A184S appeared suboptimal for multiplex imaging of longer or higher-frequency AP trains. To enable fast and high-resolution imaging of presynaptic Ca^2+^ and glutamate release in individual axonal boutons we therefore focused on the combination of Cal-590 and iGluSnFR.A184V fluorescence signals acquired in a multiplex tornado mode.

**Figure 2.**
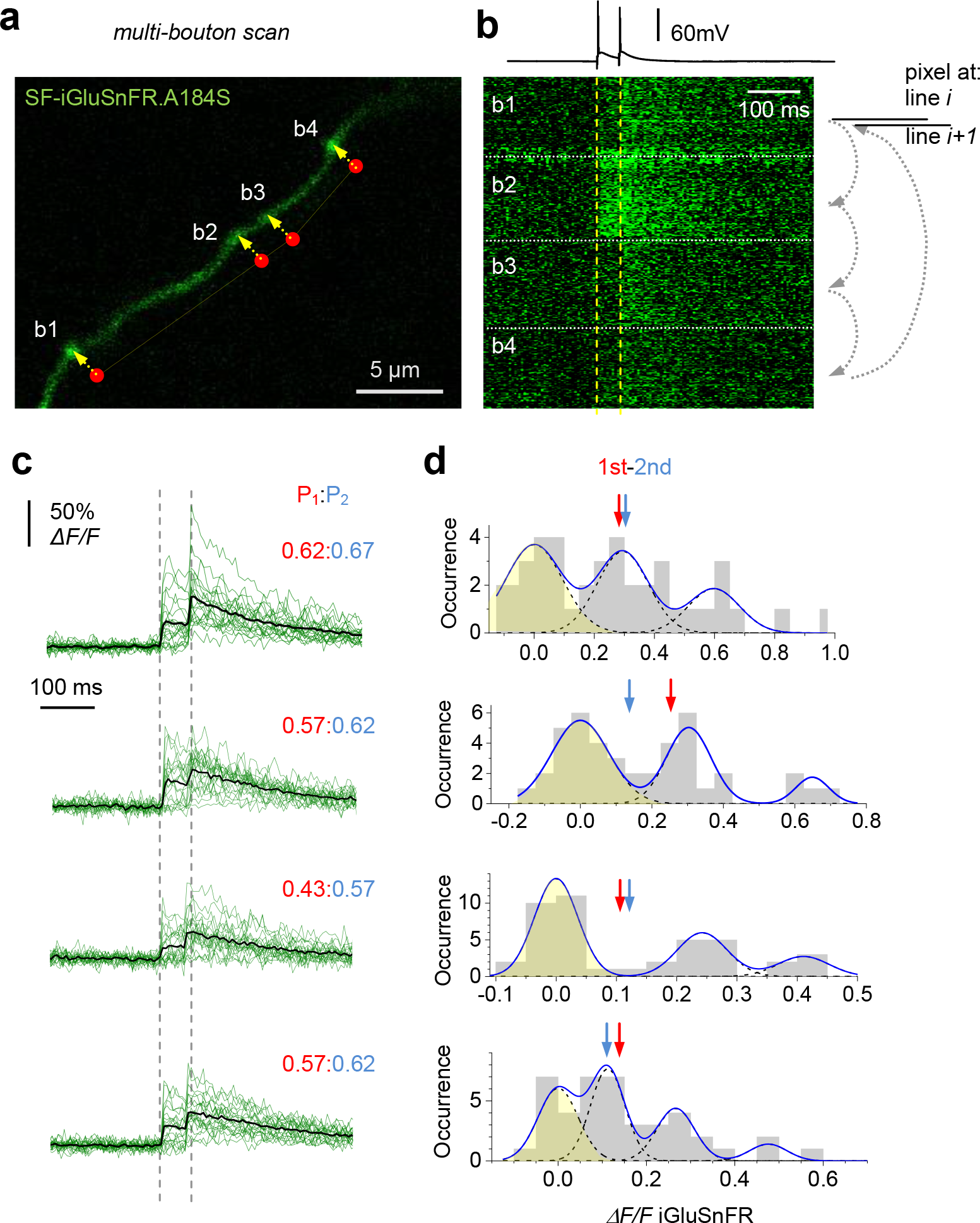
Multi-synapse imaging of quantal glutamate release with iGluSnFR.A184S. **(a)** CA3 pyramidal cell axon fragment in area CA1 showing four presynaptic boutons (b1-b4); the scanning dwell points in the bouton centres (red dots, dwell-delay time ~1.5 ms per bouton) and laser scan trajectory (dotted yellow line) illustrated. **(b)** A pseudo-linescan image of SF-iGluSnFR.A184S signals recorded simultaneously (one sweep example) at four boutons shown in **a** as indicated, during somatic generation of two APs 50 ms apart (top trace, current clamp). Arrow diagram relates displayed pixels to the scanning cycle: pixels at each displayed *i*th line are recorded sequentially among boutons (small arrows), and the next cycle fills the (*i+1*)th line of the display. Thus, a pseudo-linescan image is generated showing brightness dynamics at individual boutons, with ~1.5 ms resolution; glutamate releases and failures can be seen. **(c)** A summary of 22 trials in the experiment shown in **a-b**; green traces, single-sweep SF-iGluSnFR.184S intensity readout at the four bouton centres; black traces, all-sweep-average; P_1_:P_2_, average probability of the first (red) and second (blue) release events. **(d)** Amplitude histograms (SF-iGluSnFR.184S *ΔF/F* signal, first and second response counts combined; pre-pulse baseline subtracted), with a semi-unconstrained multi-Gaussian fit (blue line, Methods) indicating peaks that correspond to estimated quantal amplitudes; the leftmost peak corresponds to zero-signal (failure; yellow shade); dotted lines, individual Gaussians; arrows, average amplitudes (including failures) of the first (red) and second (blue) glutamate responses.

### Simultaneous monitoring of quantal glutamate release and presynaptic Ca^2+^ dynamics

Equipped with the Cal-590 FLIM calibration, we set out to simultaneously monitor glutamate release and presynaptic Ca^2+^ at axonal boutons scanned sequentially in pyramidal cells expressing SF-iGluSnFR.A184V and dialysed whole-cell with Cal-590 (Fig. 3a). To explore basal presynaptic function and its short-term plasticity, we recorded fluorescent responses, in tornado mode, to four APs evoked at 20 Hz, a pattern of activation that falls well within the range documented for CA3-CA1 connections *in vivo* ^21^. The signals were fully chromatically separated (Fig. 3b), with the SF-iGluSnFR.A184V (green) channel readily reporting individual release events in single-sweep traces, with high resolution and excellent signal-to-noise ratio (Fig. 3c-d). The Cal-590 (red) channel showed similarly high signal-to-noise ratios, and the cumulative photon count appeared sufficient to monitor the average kinetics of presynaptic [Ca^2+^] (Fig. 3e-f; note that the Ca^2+^ fluorescence readouts reflect Ca^2+^ levels that are volume-equilibrated on the 2-5 ms scale).

**Figure 3.**
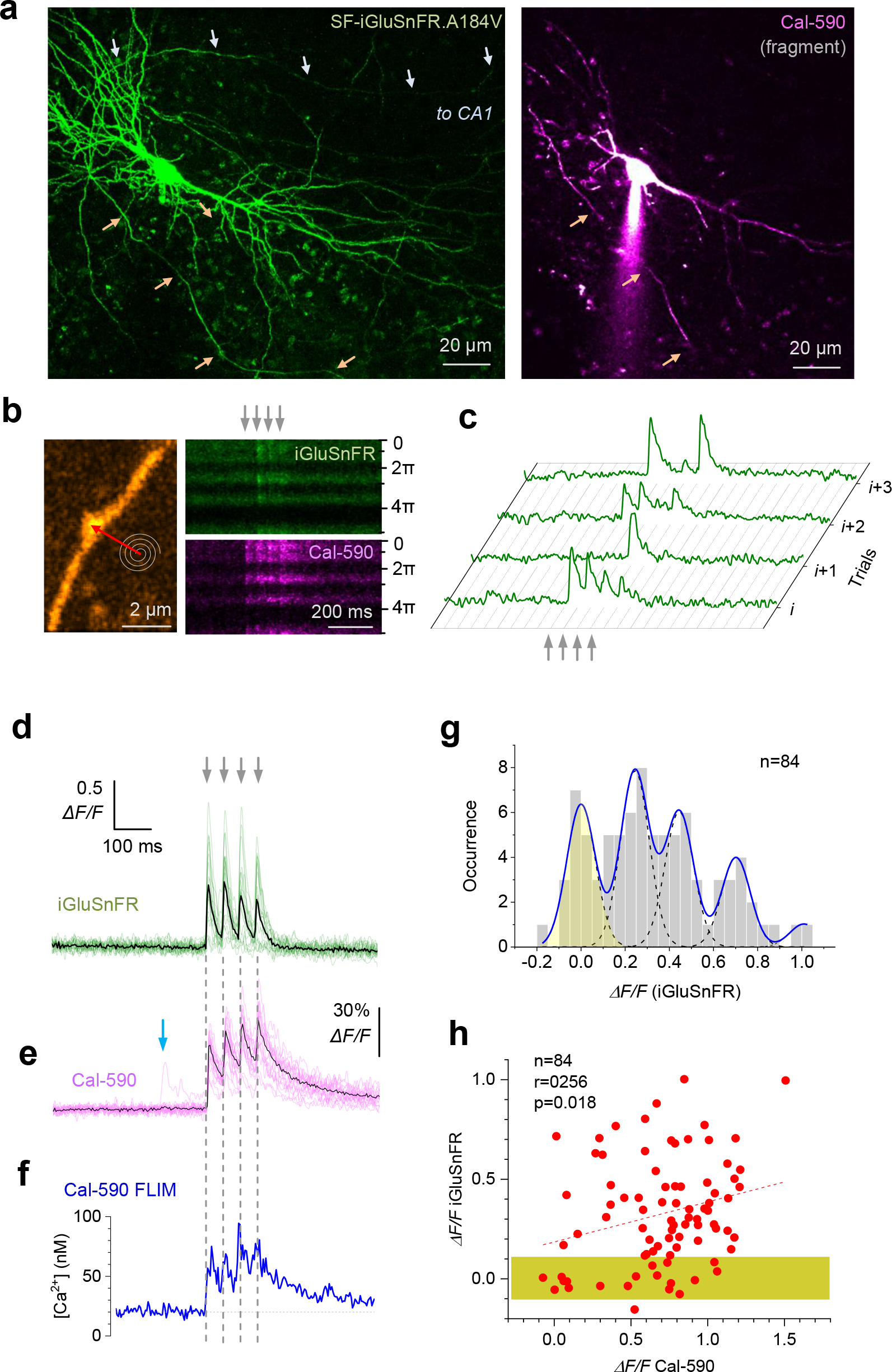
Multiplex imaging of quantal glutamate release and presynaptic Ca^2+^ dynamics. **(a)** CA3 pyramidal cell, organotypic slice. *Left:* green emission channel, biolistic transfection with SF-iGluSnFR.A184V (ref. ^17^; Methods); arrows (white/orange), two main axonal branches. Planar projection, ~60 μm deep image *z*-stack, λ_x_^2p^ =910 nm. *Right*, red emission channel (fragment of a ~20 μm *z*-stack projection), after whole-cell patch and dialysis with Cal-590 (300 μM); patch pipette is seen. **(b)** Functional imaging of an individual axonal bouton (traced into CA1 from one CA3 pyramidal cell, as detailed earlier ^8, 44^; also see Fig. 4 below): *left panels*, morphological image (iGluSnFR green channel); spiral and red arrow, tornado scan positions; *right panels*, examples of tornado line-scans (spiral rotation angle 0-5π reflects 2.5 concentric spiral circles, 2π radian/360° each) during four somatically evoked APs (20 Hz, grey arrows); shown in SF-iGluSnFR.A184V (green) and Cal-590 (magenta) channels, as indicated; note glutamate release failures in the iGluSnFR channel. **(c)** Four characteristic sequential one-sweep recordings (iGluSnFR channel) depicting individual quantal releases and failures, as indicated. **(d)** Summary of glutamate release kinetics (SF-iGluSnFR.A184V channel) documented in 20 sequential trials 1 min apart (green, individual trials; black, average trace) in bouton shown in b. **(e)** Cal-590 fluorescence intensity signal *ΔF/F* (red channel) recorded simultaneously in the test shown in **d** (magenta, individual trials; black, average trace); blue arrow, a spontaneous Ca^2+^ entry event in one trial, which is not accompanied by glutamate release (see **d**). **(f)** Dynamics of free presynaptic [Ca^2+^] averaged over 20 trials: signal readout with Cal-590 FLIM (normalised photon count), converted to [Ca^2+^]. Note that these data reflect free ion concentration volume-equilibrated and time-averaged over 5-10 ms. **(g)** Amplitude histograms (SF-iGluSnFR.184V *ΔF/F* signal, 1-4 response counts combined; pre-pulse baseline subtracted), with a semi-unconstrained multi-Gaussian fit (blue line, Methods) indicating peaks that correspond to estimated quantal amplitudes; the leftmost peak corresponds to zero-signal (failure; yellow shade); the analyses estimate noise cut-off (~50% chance of real signal) at ~0.12 *ΔF/F*. **(h)** Glutamate release signal (SF-iGluSnFR.184V *ΔF/F*) plotted against *ΔF/F* Cal-590 signal (both recorded for 1-4 *ΔF/F* Cal-590 signal responses, with 8 ms pre-AP baselines subtracted); dotted line, linear regression (r, Pearson’s correlation; p, regression slope significance); yellow shade, Gaussian noise (failure response) cut-off, ~95% confidence interval.

In this one-bouton example (Fig. 3b), the SF-iGluSnFR.A184V signal amplitude histogram analysed with multi-peak fitting (Methods; see above) revealed quantal numbers and sizes, suggesting up to four quanta per release (Fig. 3g). Importantly, the amplitudes of glutamate-sensitive responses (AP-evoked *ΔF/F* increments, measured with the ~8 ms baseline subtracted) were positively correlated with the corresponding AP-evoked [Ca^2+^] increments (Cal-590 *ΔF/F*), across the sample (Fig. 3h). This observation indicates that release probability at this synapse depends on the trial-by-trial fluctuations of AP-evoked Ca^2+^ entry.

One important consideration in the present context is that Ca^2+^ indicators such as Cal-590, by buffering intracellular Ca^2+^, could interfere with the endogenous Ca^2+^ dynamics and thus neurotransmitter release properties (and possibly with the FLIM readout conditions). To gauge these potential concomitants, we first imaged a sub-set (n = 7 cells) of axonal boutons at relatively short times after whole-cell break-in, when Cal-590 had not yet equilibrated throughout the axon. Thus, during 20-22 recording trials (1 min apart) the axonal Cal-590 concentration, hence local Ca^2+^ buffering capacity and its related effects, continued to rise, up to 2-3-fold, as monitored using total photon count of Cal-590 emission (Supplementary Fig. 4a). Remarkably, the continued Cal-590 concentration rise had no effect on resting [Ca^2+^] or the spike-evoked Ca^2+^ entry measured using Cal-590 FLIM readout, which remained perfectly stable (Supplementary Fig. 4b-c). This suggested no concomitant effects of Cal-590 fluctuations on FLIM readout, which was also in line with the tolerance of Ca^2+^ homeostasis in neuronal processes to moderate levels of Ca^2+^ buffering, as shown earlier ^12^. Furthermore, when we compared axonal boutons between iGluSnFR-expressing CA3 pyramidal cells loaded, or not loaded, with Cal-590, the average detected release probabilities were indistinguishable (Supplementary Fig. 4d). The latter observation suggests that Cal-590 loading has little overall effect on synaptic release lending further support to the physiological relevance of the present method (see Discussion for detail).

### Glutamate release depends on resting presynaptic Ca^2+^ and on AP-evoked presynaptic Ca^2+^ rises

To further understand the relationship between presynaptic Ca^2+^ dynamics and glutamate release properties further, we carried out experiments involving multiplex imaging of Cal-590 and SF-iGluSnFR.A184V (as illustrated in Fig. 3) in multiple cells recording from individual or multiple axonal boutons (Fig. 4a). Thus we were able to analyse hundreds of release events coupled with the Cal-590 FLIM readout for presynaptic [Ca^2+^]. These data revealed that the amount of released glutamate, evoked by either a single AP or a four-AP train, depended on presynaptic resting [Ca^2+^] (Fig. 4b-c). Furthermore, fluctuations in the AP-evoked presynaptic [Ca^2+^] increment were also linearly related to glutamate release efficacy (Fig. 4d), as was suggested by the intensity recordings (Fig. 3h). These observations thus indicate that basal fluctuations in release probability at small central synapses do follow fluctuations in presynaptic [Ca^2+^] rather than occurring purely independently.

**Figure 4.**
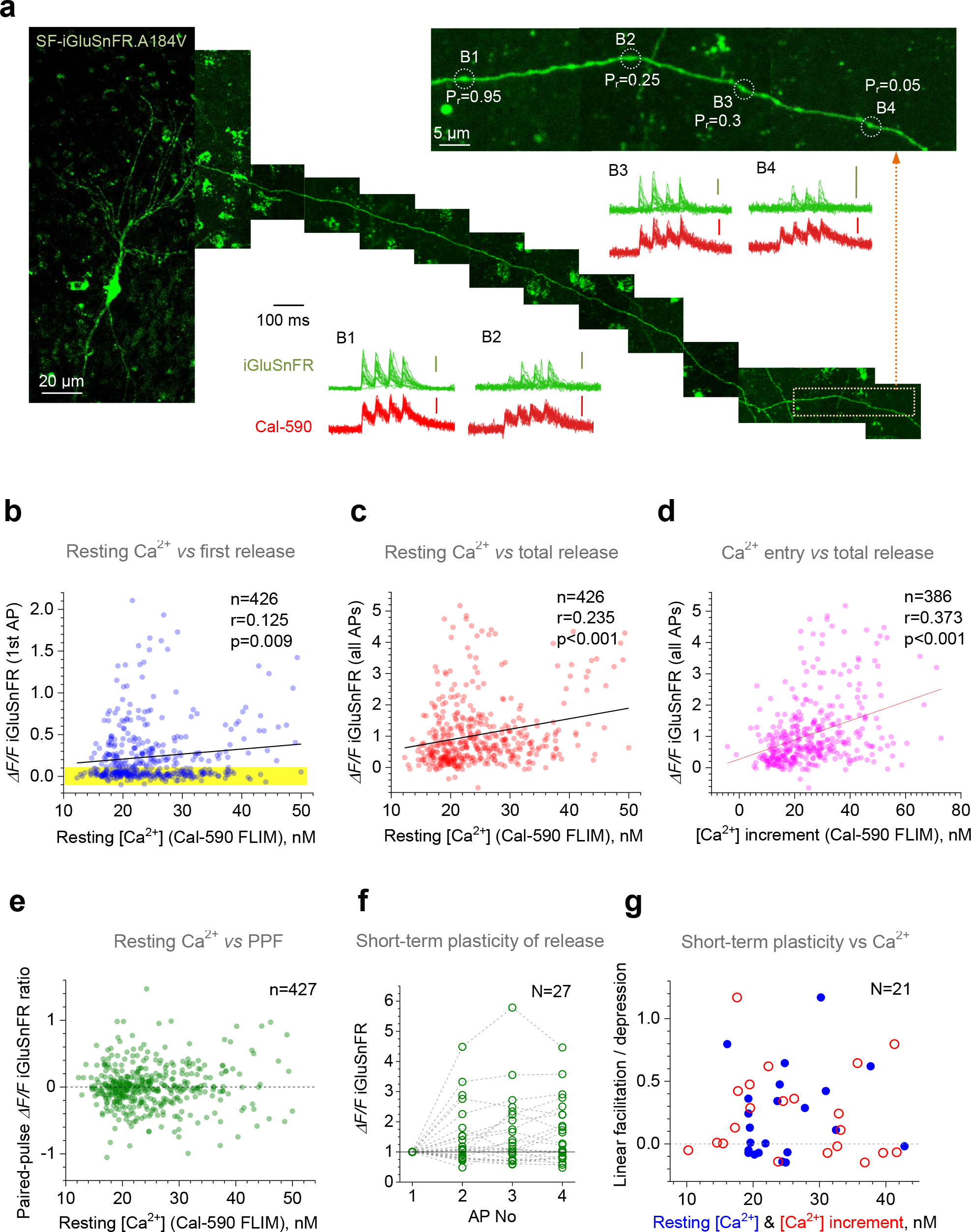
Resting presynaptic Ca^2+^ and evoked Ca^2+^ entry both control neurotransmitter release potency but not short-term facilitation at excitatory synapses. **(a)** Image diagram, CA3 pyramidal cell with its axon traced into area CA1 (example; iGluSnFR channel; collage, 2D-projections of 10-15 μm z-stacks); inset, zoomed in axonal fragment (orange dotted rectangle) depicting four recorded axonal boutons B1-B4 (P_r_, average release probability); scanning mode: tornado linescan. Traces, summary recordings of glutamate release (green) and Ca^2+^ dynamics (red), as indicated; scale bars, 0.5*ΔF/F* (green, SF-iGluSnFR.A184V) and 0.15*ΔF/F* (red, Cal-590). **(b)** Glutamate release (*ΔF/F* iGluSnFR signal) upon first AP, plotted against resting [Ca^2+^] (Cal-590 FLIM readout, averaged over 100 ms pre-pulse). Solid line, linear regression (r, Pearson’s correlation; p, regression slope significance; n, number of events, N = 26 axonal boutons recorded); yellow shade, Gaussian noise (failure response) cut-off, ~95% confidence interval. (**c**) Cumulative glutamate release upon four APs (*ΔF/F* iGluSnFR signals summed, with pre-AP 8 ms baselines subtracted) plotted against presynaptic resting [Ca^2+^] (Cal-590 FLIM readout); other notations as in **b**. (**d**) Cumulative glutamate release upon four APs (as in **c**), plotted against cumulative [Ca^2+^] increment (Cal-590 FLIM readout, during four APs); other notations as in **b** (N= 24 boutons recorded). (**e**) Paired-pulse facilitation (PPF, ratio between 2nd and 1st *ΔF/F* iGluSnFR signals, with pre-AP 8 ms baselines subtracted) plotted against presynaptic resting [Ca^2+^]. Other notations as in **b**. (**f**) Amplitude of *ΔF/F* iGluSnFR signal evoked by 1-4 APs, relative to the 1st *ΔF/F* signal, plotted against the AP number (1-4; N, number of recorded boutons). (**g**) Short-term facilitation / depression of AP burst-evoked glutamate release (calculated linear regression slope over 1-4 *ΔF/F* iGluSnFR signal change shown in **f**) plotted against presynaptic resting [Ca^2+^] (blue dots) and AP-evoked [Ca^2+^] increment (red circles).

At the same time, neither the resting Ca^2+^ level nor the variability in evoked Ca^2+^ entry had any detectable effect on the paired-pulse ratios, or short-term facilitation (14/21 synapses) or depression (7/21 synapses) shown during the four-AP train (Fig. 4e-g).

### Nanoscopic geometry of Ca^2+^ and glutamate imaging in axonal boutons

Action potentials arriving at the presynaptic bouton drive open local Ca^2+^ channels: the ensuing rapid Ca^2+^ entry triggers neurotransmitter release, in a probabilistic fashion. The effective distance between the (predominant) Ca^2+^ entry site and the release triggering site (SNARE protein complex) is considered a key factor in determining release probability hence synaptic efficacy ^15, 16, 22^. What this typical distance is, whether it changes in a use-dependent manner, whether the loci of presynaptic Ca^2+^ entry and glutamate release both move laterally in the presynaptic membrane, and whether different modes of release (spontaneous as opposed to evoked) correspond to different loci remain highly debated issues.

To understand whether these issues could be approached by multiplex imaging in the current context, we looked at the nanoscopic features of the optical settings employed. Here, individual axonal boutons express a membrane-bound (green) glutamate sensor and a cytosolic (red) Ca^2+^ indicator (Fig. 5a). The inherent blur in the imaged bouton is due to the fact that excitation and emission follow the ‘probability cloud’ set by the system’s point-spread function (PSF), which is determined in large part by the optical diffraction limit. In a well-tuned two-photon excitation microscope, the PSF is expected to be in the range of 0.2-0.3 μm in the (focal, *x-y*) plane of view, and 0.8-1.2 μm in the *z* direction, although the latter is effectively larger due to focus fluctuations. Thus, a spiral (tornado) laser scan of an axonal bouton provides roughly uniform excitation, with optical resolution of 0.2-0.3 μm in the focal plane, while integrating the fluorometric signal in the *z* direction (Fig. 5b). In other words, the imaged bouton in the current settings reflects its *z*-axis projection onto the *x-y* (focal) plane (Fig. 5c).

**Figure 5.**
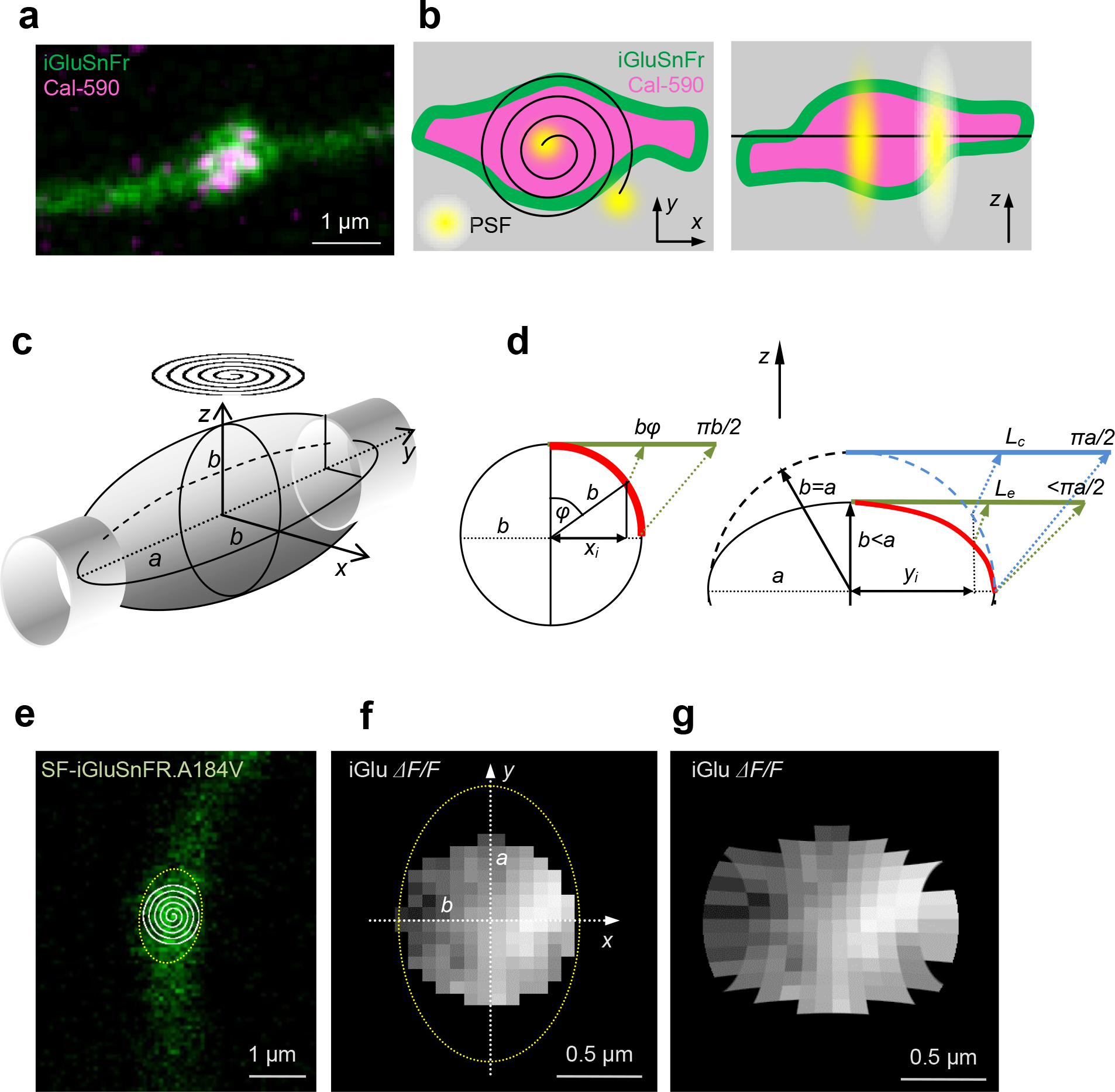
iGluSnFR imaging of presynaptic boutons reveals hotspots and spread of released glutamate. **(a)** An example of a CA3-CA1 axonal bouton, a merged-two-channel image (SF-iGluSnFR.A184V green, Cal-590 magenta). **(b)** Axonal bouton diagram illustrating imaging (laser scanning) settings, in the *x-y* (left) and *z* (right) plane, with membrane-bound expression of iGluSnFR (green) and cytosolic Cal-590 (magenta); black spiral (left) and straight line (right), approximate tornado linescan trajectory; yellow spots illustrate typical point-spread function (PSF): note that the *x-y* plane image represents, in large part, *z*-projection of the bouton. **(c)** Diagram, typical experimental arrangement, with round tornado linescan (spiral) over an ellipsoidal axonal bouton (*a* and *b*, major and minor axes, respectively, of the elliptical *x-y* projection for the rotational ellipsoid). **(d)** Trigonometry diagrams explaining geodesic corrections for distances on curved surfaces that are projected onto the (focal) *x-y* plane. *Left*, for a circle or cylinder of radius *b*, the projected distance *x*_*i*_ from the centre corresponds to the curvilinear (geodesic) distance *bφ* where angle *φ* = arcsin (*x*_*i*_/*b*) (green segment depicts geodesic distance ‘straightened’ in plane of view); thus, the projected distance from the centre to the edge, *b*, corresponds to the geodesic distance π*b*/2. *Right*, for an elliptical section with major and minor axes *a* and *b*, respectively, the projected distance *x*_*i*_ from the centre corresponds to the geodesic distance (*L_e_*, green segment), which is smaller than that for a circular correction (*L*_*c*_, blue segment). The difference between *L*_*e*_ and *L*_*c*_ depends on the expected *a* / *b* ratio (see text and Supplementary Fig. 4d-f for detail). **(e)** An example of a recorded bouton (SF-iGluSnFR.A184V channel), with a tornado scan shown; dotted oval, an estimated outline of the axonal bouton projection. **(f)** Characteristic heat map of the *ΔF/F* SF-iGluSnFR.A184V fluorescence signal generated by an AP-evoked glutamate discharge (one-bouton example; imaging arrangement as in **e**, circular shape follows the tornado scan); average of 21 trials 1 min apart; dotted oval, bouton outline as shown in **e**; image projected on to the focal *x-y* plane. **(g)** Heat map of *ΔF/F* iGluSnFR as in **f** corrected for geodesic distances as opposed to the projected (visible) distances, in accord with the geometry of the bouton outline and the tornado scan position.

Synaptic vesicle release generates a steep diffusion gradient, with the extracellular glutamate concentrations falling orders of magnitudes over tens of nanometres from the release site ^15, 16, 22–24^. Thus, the respective fluorescent signal, such as one from SF-iGluSnFR.A184V, is expected to show distinct nanoscopic hotspots. However, because this signal originates from near the 3D bouton surface, it is important to understand how it is projected onto the microscope plane of view. The bouton projection that is registered by the circular tornado scan (Fig. 5c) underestimates curvilinear (geodesic) distances on the bouton surface. Measuring the profiles of 26 recorded axonal boutons suggested that most could be reasonably approximated by rotational ellipsoids (Supplementary Fig. 5a-b), with the visible rotation radius of 0.89 ± 0.04 μm and the major axis radius (along the axon trajectory) of 1.18 ± 0.04 μm (Supplementary Fig. 5c).

Geodesic correction for curvilinear distances on the spherical surface of radius *b* is straightforward: the projected distance *x*_*i*_ from the profile centre will correspond to the curvilinear geodesic distance of *b* arcsin (*x*_*i*_/*b*) (Fig. 5d, left). Obtaining the exact correction for an elliptical surface (Fig. 5d, right) could be much more complicated. However, in our experimental sample the average ratio of major and minor ellipsoid axes *a / b* for axonal boutons is 1.35 ± 0.04 (Supplementary Fig. 5c). In the conservative case of *a / b* = 1.5, one can show that the difference between the corrected distances for the ellipsoidal surface (radii *a* and *b*) and the spherical surface (radius *a*) is unlikely to exceed ~5% (Supplementary Fig. 5e-f). Thus the visible planar map of the oval (ellipsoidal) bouton registered with a circular tornado scan for the SF-iGluSnFR.A184V *ΔF/F* signal (Fig. 5e-f) will stretch the visible lengths in the directions *x* and *y* by factors *b* arcsin (*x*_*i*_/*b*) and *a* arcsin (*y*_*i*_/*a*), respectively, to obtain a corrected geodesic surface map (Fig. 5g).

In addition to the geodesic distance correction, fluorescence intensity signal collected in a planar projection is overestimated towards the edge of spherical or ellipsoidal shapes (Supplementary Fig. 5g, inset). In the present study we focused on the ‘ratiometric’ *ΔF/F* iGluSnFR.A184V signal, which should effectively nullify this bias. However, because this correction could prove useful in other case, we have detailed its theory and tested it using a micro-vesicle with a fluorescent shell (Supplementary Fig. 5h-i).

### Sub-microscopic localisation of presynaptic glutamate release and Ca^2+^ entry hotspots

In 23 individual presynaptic boutons, tornado-scan heat maps (averaged over 20-22 consecutive trials, 1 min apart, in each bouton) revealed clear hotspots for the *ΔF/F* SF-iGluSnFR.A184V signal, with most synapses (20/23) showing one and the rest (3/23) showing two separate peaks (Figs. 5f, 6a). The fact that the signal averaging over multiple trials reveals individual hotspots suggests that trial-by-trial glutamate release is constrained to a relatively narrow area of the bouton membrane. We next used geodesic correction as described above to establish the spatial spread of the iGluSnFR signal across the bouton surface. The average brightness profile (n = 23 boutons) normalised to the peak value was well fitted by a single exponent with a spatial decay constant of 0.547 ± 0.016 μm (n = 23 boutons; Fig. 6b). This was in excellent agreement with biophysical studies predicting negligible glutamate escape beyond 0.5-0.7 μm at hippocampal synapses ^25^, mainly due to the powerful, high-affinity uptake by the surrounding astroglia ^26^.

**Figure 6.**
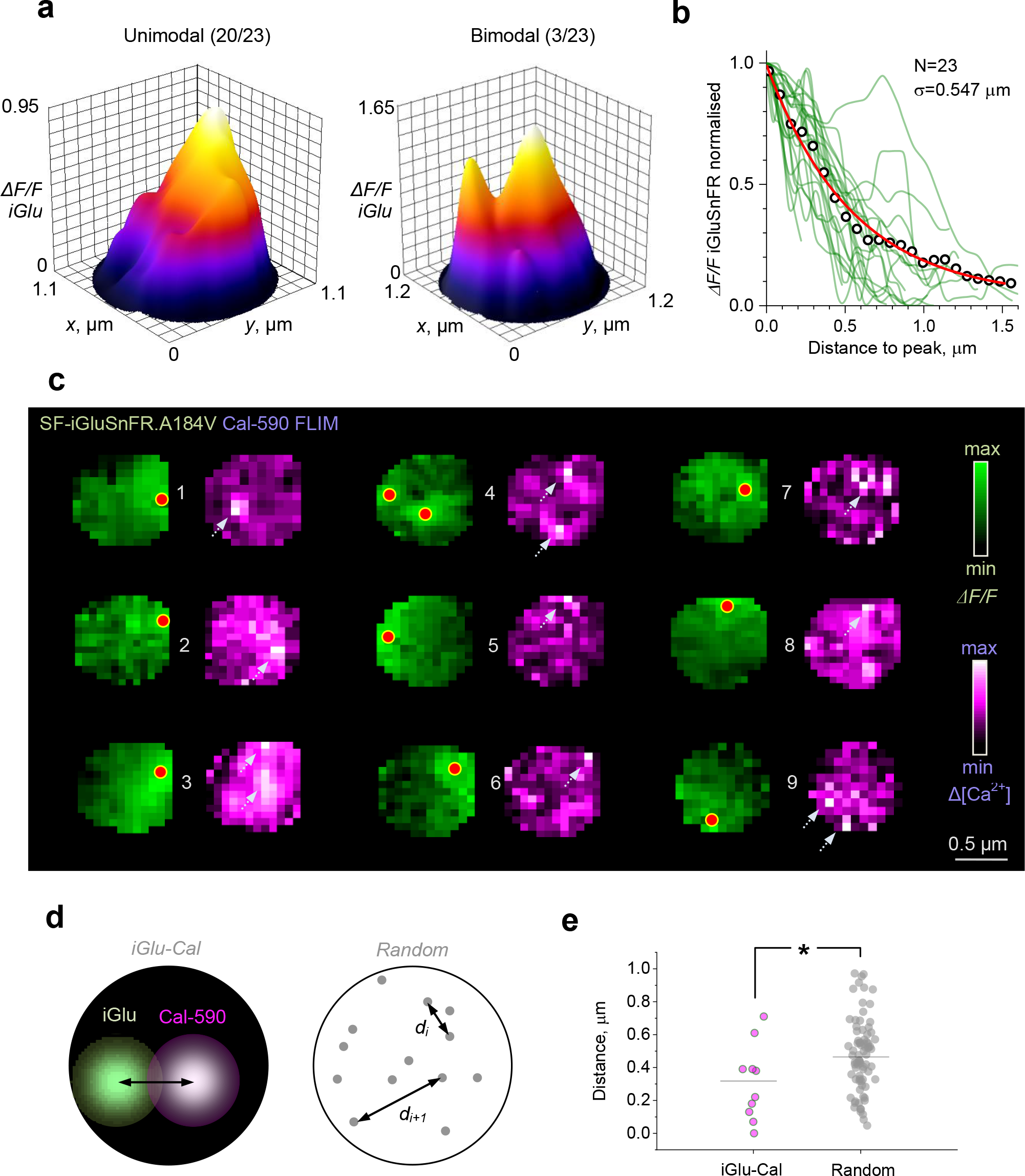
Evaluating sub-microscopic glutamate signal spread and co-localisation of glutamate release and presynaptic Ca^2+^ entry in individual presynaptic boutons. **(a)** Examples of unimodal (left, 20/23 boutons) and bi-modal (right, 3/23 boutons) spatial profiles of the iGluSnFR signal upon glutamate release, as seen in the focal plane; averaging spatial filtering (~100 nm range) applied; peaks point to the most likely release site location. **(b)** Spatial decay of the fluorescence signal (*ΔF/F* SF-iGluSnFR.A184V) measured in geodesic-corrected heat maps (as in Fig. 5g) for N=23 individual boutons (green lines); circles, average; solid red line, best-fit exponent: spatial decay constant σ = 0.547 ± 0.016 μm (mean ± SE). **(c)** Examples of simultaneously recorded heat maps showing glutamate signal (green, *ΔF/F* iGluSnFR) and [Ca^2+^] increment profiles (magenta, Cal-590 FLIM readout upon four APs, magenta); red dots, estimated glutamate release site locations; arrows, consistent hotspots of Cal-590 [Ca^2+^] transients. **(d)** Diagrams illustrating the measurement of distances between recorded hotspots of glutamate and Ca^2+^ signals, as shown in **c** (left), and between points randomly scattered within a similar circular area (right). **(e)** The average distance between recorded hotspots of glutamate and Ca^2+^ signals (0.308 ±0.073 μm, n = 10; grey dots) is significantly lower than that between randomly scattered points (0.474 ± 0.025, n = 80, orange dots; p < 0.028).

In these experiments, multiplex imaging of Cal-590 and iGluSnFR.A184V provided the opportunity to assess spatial co-localisation of the strongest Ca^2+^ entry and the glutamate release site (Fig. 6c; Methods). Whilst glutamate release heat maps consistently displayed well-identified hotspots, the Cal-590 FLIM readout signal averaged over 20-22 trials in individual boutons showed lower signal-to-noise ratios compared to the iGlu readout (Fig. 6c). This was likely due to a combination of factors including a suboptimal total photon count, variable pattern of intra-bouton Ca^2+^ channel activation signal in individual trials, or even their lateral movement between trials (see Discussion). Nonetheless, clear Ca^2+^ entry hotspots (>2SD of the background noise) were detected in at least nine synaptic boutons, thus enabling the assessment of their co-localisation with the glutamate release site (Fig. 6c).

Firstly, in the data sample only 4 out of 9 synapses appeared to show Ca^2+^ entry-glutamate release co-localisation in the 0-200 nm range (pairs 6,7,8,9 in Fig. 6c; because Ca^2+^ hotspots could occur within the bouton volume, these maps were not corrected for geodesic distances). To test whether the recorded juxtaposition pattern occurred purely by chance, we ran a straightforward statistical test in which the experimental scatter of distances between Ca^2+^ and glutamate hotspots was compared with the scatter of distances among arbitrary pairs of points randomly scattered over a similar circular area (Fig. 6d). The test revealed that the experimental distances were significantly lower than the simulated ones (Fig. 6e), suggesting that Ca^2+^ entry occurs significantly closer to the site of glutamate release than it would be expected purely by chance. Functional implications of this observation remain an open and intriguing question.

## DISCUSSION

### FLIM monitoring of low presynaptic Ca^2+^ with Cal-590

In this study we have established that the fluorescence lifetime of the red-shifted Ca^2+^ indicator Cal-590 is sensitive to [Ca^2+^] in the 10-200 nM range, with an optimal two-photon excitation wavelength of ~910 nm. We have calibrated this dependence using the normalised photon counting protocol described earlier ^12, 13^ and thus been able to obtain direct readout of [Ca^2+^] dynamics in neuronal axons using whole-cell dialysis of CA3 pyramidal cells with Cal-590 in organotypic brain slice preparations. The knowledge about low basal [Ca^2+^] and its changes in neuronal axons is fundamentally important for understanding the machinery of neurotransmitter release. This is because free [Ca^2+^] level reflects equilibrium between local Ca^2+^ entry, removal, and binding-unbinding with local Ca^2+^ buffers. Thus, changes in low [Ca^2+^] not only report a shifted equilibrium, they directly alter the availability (and thus buffering capacity) of Ca^2+^-free endogenous Ca^2+^ buffers ^27–29^. As the bound-to-free ratio for presynaptic Ca^2+^ varies between 100-1000-fold ^18, 22, 30^, equilibrated basal Ca^2+^ must contribute strongly to the spatiotemporal integration of intracellular Ca^2+^ ‘hotspots’ hence the triggering of release ^15, 16^. Strong Ca^2+^ buffering can also explain why the range of presynaptic [Ca^2+^] values we have detected is remarkably low, representing only a few free Ca^2+^ ions per cubic micron. It is important to understand that these are never the same few molecules, as the local bound Ca^2+^ is in a continued and rapid exchange with free cytosol Ca^2+^.

In addition to high [Ca^2+^] sensitivity in the nanomolar range, at least in the case of OGB-1 or Cal-590, a critical technical advantage of the FLIM readout is that it does not depend on the dye amount or concentration, light scattering, focus fluctuations, and other concomitants of live imaging ^12, 13^. The latter is particularly important for imaging in freely-moving animals. Having a relatively more complicated procedure to record and analyse FLIM recordings compared to standard fluorometric monitoring ^12, 13^ appears a small price to pay for such advantages.

### Intracellular Ca^2+^ buffering by Cal-590

Because Ca^2+^ indicators are by definition Ca^2+^ buffers they could affect intracellular Ca^2+^ homeostasis. Our recent study has shown, however, that moderate increases in the intracellular concentration of OGB-1 has little influence on the cell-wide landscape of low basal Ca^2+^ in principal neurons ^12^. This is because inside cells the steady-state, equilibrated [Ca^2+^] level depends on the action of Ca^2+^ sources and sinks, which in turn are controlled by active [Ca^2+^] sensing mechanisms rather than by ‘passive’ Ca^2+^ buffers ^28^. However, the effect of Ca^2+^ indicators on rapid presynaptic Ca^2+^ dynamics and release probability could be significant, as we demonstrated previously ^18, 31^. Cal-590 has a relatively low affinity (*K*_*d*_ ~ 560 nM) compared to OGB-1 (*K*_*d*_ ~ 170-260 nM) ^11, 32^, and should therefore have an even lower Ca^2+^ buffering effect. Intriguingly, we found no significant effect of axon loading with Cal-590 on the average release probability. Whilst this speaks in favour of the method, the physiological implications of this observation, in terms of the presynaptic mechanisms involved, require a separate study. Reassuringly, it was earlier reported that bolus-loading application of Cal-590 was fully compatible with intense network spiking activity in the visual cortex *in vivo* ^14^. Furthermore, Ca^2+^ indicators with higher affinity, such as OGB-1 or some genetically-encoded Ca^2+^ indicators, have been widely used to monitor neural network activity in freely-moving animals, without apparent behavioural implications ^33, 34^. In any case, a more systematic comparison of release probabilities among axons loaded with Cal-590 at different concentrations, within the same synaptic circuitry, are required to quantitatively assess potential subtle effects of the dye.

### Multiplex imaging reveals basic relationships between presynaptic Ca^2+^ and glutamate release at small central synapses

Enabling simultaneous readout of FLIM-enabled presynaptic Ca^2+^ and quantal glutamate release at small central synapses *in situ* is important for at least two reasons. Firstly, it enables a monitoring regime of release probability that is insensitive (or only weakly sensitive) to the fluorescence emission intensity registered in organised brain tissue. Thus, the common concomitants of live imaging *in situ* or *in vivo*, such as focus drift, photobleaching, or physiological changes in tissue light scattering or absorption should have only a limited effect on the release probability readout with the present multiplex approach. Furthermore, axonal Ca^2+^ monitoring *in vivo* is essential for presynaptic spike detection: the efficiency of (bolus-applied) Cal-590 in registering APs in deep brain imaging settings has recently been demonstrated ^14^. Thus, comparing the occurrence of spikes (Ca^2+^ channel) with the statistics of stochastic glutamate release events (iGluSnFR channel) at individual axonal boutons should provide a quantitative gauge of synaptic transmission fidelity, and its use-dependent changes, in such experiments.

Secondly, the method provides insights into the mechanistic relationship between Ca^2+^ entry and release probability, and how this relationship changes during use-dependent synaptic plasticity. To address such fundamental questions, classical studies in giant (Calyx) synapse preparations combined presynaptic Ca^2+^ imaging with patch-clamp recordings in pre- and/or postsynaptic structures, which were large enough to allow access ^9, 10, 35^. Whilst such studies have been pivotal for understanding the basics of Ca^2+^-dependent release machinery, to what degree their conclusions could be extrapolated to diverse neural circuits, in different brain regions, remains poorly understood. In this context, it is particularly important to have reliable readout of low basal Ca^2+^, which directly affects the saturation level hence local availability of Ca^2+^-free buffers ^10^. The present study reveals that fluctuations in basal presynaptic [Ca^2+^] as well as in the AP-evoked Ca^2+^ entry affect release efficacy. The role of presynaptic depolarisation-induced rises in basal [Ca^2+^] for neurotransmitter release has previously been shown in giant calyceal synapses^36^. Our data suggest that the basal [Ca^2+^] - release efficacy relationship is maintained in small central synapses, during quiescent conditions in situ. Whether the fluctuations in presynaptic resting [Ca^2+^] occur purely stochastically or are driven by the external (network) factors remains to be ascertained. Similarly, the observation that fluctuations in AP-evoked Ca^2+^ entry are also correlated makes such fluctuations a significant contributor to the stochastic nature of neurotransmitter release, in addition to other presynaptic mechanisms.

The kinetics of the glutamate sensor variant SF-iGluSnFR.A184S (as well as the kinetics of Cal-590) used in this study are well suited to monitor release probability at multiple synapses supplied by the same axon. Thus, the method enables real-time readout of heterogeneous presynaptic identities, their possible collective or cooperative features, and their plasticity within an identified synaptic circuit. On a technical note, with the present method, we used the basal iGluSnFR fluorescence signal, under sparse expression of the indicator, to trace individual cell axons. Clearly, this axonal tracing approach could be adapted to the use of other appropriate dyes and genetically encoded indicators (such as a red-shifted version of iGluSnFR ^37^ or sensors for other neurotransmitters), or other sparse transfection methods, to enable simultaneous multi-target *in vivo* functional imaging. In addition to transfection of indicators, co-transfection with RNAi or CRISPR costructs should offer a method to assess the molecular signalling mechanisms underlying glutamate release and short-term plasticity ^38^.

### Nanoscopic localisation of presynaptic glutamate release and Ca^2+^ entry

Release of glutamate involves membrane fusion of a 40 nm synaptic vesicle, generating a sharp neurotransmitter diffusion gradient inside and outside the cleft ^24, 39^. Similarly, Ca^2+^ entering presynaptic boutons through a cluster of voltage-gated Ca^2+^ channels dissipates orders of magnitude at nanoscopic distances from the channel mouth ^15, 16, 22^. Thus, when the respective fluorescent indicators are present they should in both cases generate a nanoscopic signal hotspot. In recorded images such hotspots will be blurred by the microscope’s PSF and the inherent experimental noise. However, taking advantage of averaging under multiple exposure, over repeated trials, we were able to localise the preferred regions of glutamate release and, in some cases, Ca^2+^ entry sites, beyond the expected diffraction limit of ~250 μm in the *x-y* plane (Fig. 6a-c). Clearly, such accuracy in the *z* direction appears more problematic: it will require specific optics that would enable registration of optical aberration (such as astigmatism) against the *z* coordinate ^40^ or otherwise a specifically modified PSF shape ^41^.

In our experiments, only some synapses displayed co-localisation of Ca^2+^ and glutamate signal hotspots on the small scale (<200 nm), although across the sample the two sites on average occurred significantly closer to one another than it would be expected from their random positioning (Fig. 6). The apparent lack of tight co-localisation lends support to the view of loose coupling between presynaptic Ca^2+^ channels and synaptic release machinery, which seems characteristic for plastic synapses ^16^ and has been suggested by biophysical models of axonal boutons at CA3-CA1 connections ^42^. The loose-coupling hypothesis would also appear consistent with the finding that trial-to-trial fluctuations in presynaptic basal Ca^2+^ or Ca^2+^ entry affect glutamate release efficacy (Fig. 4b-d): in such cases release should depend on the variable cooperative action of multiple Ca^2+^ channels rather than on one channel opening ^15^. Furthermore, a large proportion of recorded boutons did not shown clear Ca^2+^ entry hotspots in the all-trial-average heat maps, suggesting that the site of Ca^2+^ entry might not be consistent from trial to trial, either because of stochastic opening among a set of presynaptic Ca^2+^ channels, due to their lateral motility ^43^, or indeed because of the fluctuating contribution from local Ca^2+^ stores ^44^.

In contrast, the average glutamate signal heat maps do show nanoscopic hotspots suggesting consistent localisation of the glutamate release site from trial to trial (Fig. 6). This appears in line with the notion that some key presynaptic proteins within the active zones (in particular Rab3-interacting molecule, RIM, associated with release machinery) closely align, across the synaptic cleft, with postsynaptic receptors, in a trans-synaptic molecular ‘nanocolumn’ ^45^: such alignment would appear functionally irrelevant with a variable release site. The present multiplex imaging approach, while enabling evidence for real-time physiological events *in situ*, provides resolution for glutamate release localisation which could also be achieved by imaging vesicle-associated pHluorins that increase fluorescence intensity upon vesicle fusion ^5^. An elegant variant of this method, termed pHuse, has used nano-localisation of individual release events in cultured cells, to detect spatially constrained glutamate release sites, at least in a group of relatively small presynaptic boutons ^45^, in line with the present observations.

## METHODS

### Organotypic slice culture preparation

Organotypic hippocampal slice cultures were prepared and grown with modifications to the interface culture method ^46^ from P6-8 Sprague-Dawley rats, in accordance with the European Commission Directive (86/609/EEC) and the United Kingdom Home Office (Scientific Procedures) Act (1986). 300 μm thick, isolated hippocampal brain slices were sectioned using a Leica VT1200S vibratome in ice-cold sterile slicing solution consisting (in mM) of Sucrose 105, NaCl 50, KCl 2.5, NaH2PO4 1.25, MgCl2 7, CaCl2 0.5, Ascorbic acid 1.3, Sodium pyruvate 3, NaHCO3 26 and Glucose 10. Following washes in culture media consisting of 50% Minimal Essential Media, 25% Horse Serum, 25% Hanks Balanced Salt solution, 0.5% L-Glutamine, 28 mM Glucose and the antibiotics penicillin (100U/ml) and streptomycin (100 μg/ml), three to four slices were transferred onto each 0.4 μm pore membrane insert (Millicell-CM, Millipore, UK), kept at 37°C in 5% CO2 and fed by medium exchange every 2-3 days for a maximum of 21 days in vitro (DIV). The slices were transferred to a microscope recording chamber with the recording ACSF solution containing (in mM): NaCl 125, NaHCO3 26, KCl 2.5, NaH2PO4 1.25, MgSO4 1.3, CaCl2 2 and glucose 16 (osmolarity 300-305 mOsm), continuously bubbled with 95% O2/5% CO2. Recordings were carried out at 33-35C, with addition of 10μM NBQX and 50μM AP5 to reduce the potential for plasticity effects influencing synaptic properties during prolonged recordings.

### Biolistic transfection of iGluSnFR variants

Second generation iGluSnFR variants SF-iGluSnFR.A184S and SF-iGluSnFR.A184V ^17^ were expressed under a synapsin promoter in CA3 pyramidal cells in organotypic slice cultures using biolistic transfection techniques adapted from manufacturer’s instructions. In brief, 6.25 mg of micron Gold micro-carriers were coated with 30μg of SF-iGluSnFR plasmid. Organotypic slice cultures at 5DIV were treated with culture media containing 5 μM Ara-C overnight to reduce glial reaction following transfection. The next day cultures were shot using the Helios gene-gun system (Bio-Rad) at 120psi. The slices were then returned to standard culture media the next day and remained for 5-10 days before experiments were carried out.

### Axon tracing and imaging of presynaptic boutons

We used a Femtonics Femto2D-FLIM imaging system, integrated with patch-clamp electrophysiology (Femtonics, Budapest) and linked on the same light path to two femtosecond pulse lasers MaiTai (SpectraPhysics-Newport) with independent shutter and intensity control. Patch pipettes were prepared with thin walled borosilicate glass capillaries (GC150-TF, Harvard apparatus) with open tip resistances 2.5-3.5 MΩ. For CA3 pyamidal cells internal solution contained (in mM) 135 potassium methanesulfonate, 10 HEPES, 10 di-Tris-Phosphocreatine, 4 MgCl2, 4 Na2-ATP, 0.4 Na-GTP (pH adjusted to 7.2 using KOH, osmolarity 290-295), and supplemented with Cal-590 (300 μM; AAT Bioquest) for FLIM imaging.

Pre-synaptic imaging was carried out using an adaptation of previously described methods for separate pre-synaptic glutamate and Ca^2+^ imaging ^8^. Cells were first identified as iGluSnFR expressing using two-photon imaging at 910 nm and patched in whole cell mode as above. Following break-in, 30-45 minutes were normally allowed for Cal-590 to equilibrate across the axonal arbour; in a sub-set of experiments, shorter post-break-in times were implemented to test release properties under the continued equilibration (concentration rise) of Cal-590 inside the axon. Axons, identified by their smooth morphology and often tortuous trajectory, were followed in frame scan mode to their targets and discrete boutons were identified by criteria previously demonstrated to reliably match synaptophysin labelled punctae ^47^.

For fast imaging of action-potential mediated iGluSnFR and Cal-590 fluorescence transients at individual boutons, a spiral shaped (tornado) scan line was placed over the bouton of interest, as described previously ^8^ and illustrated in detail in Fig. 5. The spiral tornado mode is a built-in beam scanning option in the Femtonics Femto2D (or Olympus FluoView FV1000) microscope imaging system, allowing direct control over its radius, position, and scanning frequency. The sampling frequency used in the present settings was ~500 Hz, with two-photon excitation at a wavelength λ_x_^2p^ = 910 nm, with the power under objective of 3-5 mW. Through the experimental trials, axonal boutons maintained stable morphological and functional features (release probability, nanomolar Ca^2+^ level, evoked Ca^2+^ entry; Supplementary Fig. 4a-c), thus providing direct functional evidence for the experiment-wise absence of photo-toxicity effects.

For simultaneous multi-bouton imaging, the scanning mode was the sequential point-scans over selected ROIs (axonal bouton centres), adjusted to a temporal resolution of ~4 ms (~250 Hz rate). The continued cycled series of point-scans over different boutons was subsequently rearranged using a MATLAB algorithm to represent ‘pseudo-linescans’ associated with individual boutons, as illustrated in Fig. 2b. Following a baseline period two or four APs, 50 ms apart, were generated using 2 ms positive voltage steps at the cell soma in whole-cell voltage clamp mode (V_m_ holding at −70mV).

### FLIM calibration for [Ca^2+^] readout and NTC method

The Cal-590 [Ca^2+^] calibration protocol was similar to the standard calibration method provided by the Invitrogen Ca^2+^ calibration buffer kit manual. The practical procedure in large part replicated the previously described FLIM calibration protocol for OGB-1 ^12, 13^. In brief, to match the Ca^2+^ buffering dynamics to that of Cal-590 more closely, the standard 10 mM chelating agent EGTA was replaced with 10 mM BAPTA, and the solution constituents were replaced with the experimental intracellular solution (see above). pH was adjusted using KOH, and the KCl concentration was adjusted accordingly, to keep ion constituents in the solution unchanged. The calculated [Ca^2+^] was therefore slightly different from the standard Invitrogen’s calibration set, and was finely adjusted using Chris Patton’s WEBMAXC program (Stanford University, http://www.stanford.edu/~cpatton/webmaxcS.htm). In the control tests, excitation wavelength was varied between 800 nm and 950 nm, and temperature was varied from 20°C to 37°C (using a Scientific Systems Design PTC03 in-line heater).

The analysis of fluorescence lifetime data using classical multi-exponential fitting has long been the bottleneck in the experimental throughput. Because the physics that could be inferred from multi-exponential fitting was outside the scope of our studies, we earlier proposed to replace such exponential fitting with a simpler and computationally more economical approach based on a ratiometric method of normalised total count (NTC) ^12, 13^. The fluorescent decay time course was first normalized to its peak value and then integrated (area under curve) over ~3 ns post-peak (Fig. 1a). The resulting value termed NTC was used throughout as an estimator for [Ca^2+^] using an appropriate calibration function: the latter was obtained through direct Cal-590 calibration for intracellular solutions of clamped [Ca^2+^], as explained above, in the microscope imaging system used for experimental measurements (Femto2D, Femtonics, Budapest). The output of Cal-590 lifetime at different clamped [Ca^2+^] values produced almost noiseless decay traces (Fig. 1a) as the photon collection time is not limited by the experimental conditions. In turn, applying the NTC method to these traces and plotting the outcome against the estimated [Ca^2+^] produced a near-perfect fit for logistic function (Fig. 1b).

### Two-photon excitation FLIM of Cal-590 *in situ*

With the scanning modes and methods described above, the line-scan data were recorded by both standard analogue integration in Femtonics MES and in TCSPC in Becker and Hickl SPCM using dual HPM-100 hybrid detectors. The two-photon laser source was a Newport-SpectraPhysics Ti:Sapphire MaiTai laser pulsing at 80 Mhz, with a pulse width of ~220 fs and a wavelength of 910 nm optimized for Cal-590 excitation.

FLIM line scan data were collected as previously described ^12, 13^ and stored as 5D-tensors (*t,x,y,z,T*) where *t* and *T* stand for the lifetime decay timing and the (global) experiment-wise timing, respectively. The corresponding off-line data sets were analysed using custom written analysis software (https://github.com/zhengkaiyu/FIMAS). For bouton-average Cal-590 FLIM measurements, the *x*, *y* and *z* data sets were summed along their respective axes. In contrast, to obtain FLIM readout maps within individual boutons, the recorded photon counts in all relevant co-ordinates (tornado scanning mode) were summed across trials (*T*) to produce count numbers sufficiently high for the reliable NTC measure.

### Quantal analyses of evoked iGluSnFR signals

Optical glutamate imaging provides a unique opportunity to monitor release activity at individual identified synapses in a non-invasive manner. Similar to the classical electrophysiological studies of synaptic transmission, in this case the quantal size and content of neurotransmitter discharges can be assessed using frequency histograms reflecting trial-to-trial fluctuations of the signal amplitude ^20^. In such histograms, amplitude fluctuations around zero (failure) responses reflect the signal noise inherent to the particular recording settings. Establishing the noise fluctuation using a straightforward unsupervised Gaussian fitting (with one free parameter, dispersion σ; and two fixed parameters, centre *x* = 0, and intersect *y* = 0) will thus constrain (approximately) the key noise parameter σ which should apply to all quantal sizes. In the present study, once σ has been constrained, we set up initial parameter values (peak centres *x*_*i*_), their margins (±15%), σ*i* margins (±15%), and employed the unsupervised Levenberg-Marquardt iteration algorithm for multi-Gaussian fitting (GaussAmp function; Origin, OriginLab). The outcome produced individual Gaussians that correspond to the peaks of quantal amplitude, and the summated distribution (Figs. 2d; 3g; Supplementary Fig. 2f).

### Nanoscopic localisation of presynaptic glutamate release and Ca^2+^ entry

In individual axonal boutons, the heat maps of iGluSnFR.A184V signal collected with tornado scans at ~500 Hz were average across individual trials along the corresponding time points in the trial. For the period AP-evoked activity the maps were plotted as *ΔF/F* signal readouts with the basal signal F calculated over the 300 ms pre-pulse interval. A similar procedure was applied to the Cal-590 FLIM readout signal, with heat maps of the baseline signal (*F*) reporting resting [Ca^2+^], and with the *ΔF/F* maps during the period of AP-evoked activity reporting AP-evoked [Ca^2+^] increments.

Spatial localisation of the glutamate release site was carried out in the average *ΔF/F* SF-iGluSnFR.A184V heat maps (collected during the period between first AP and 15 ms after the fourth AP). This, the maps represent averaged signals between 2 ms before and 10 ms after the AP-induced peaks, thus excluding the values in troughs of the waveform. Because the registered glutamate signal hotspots were generally asymmetrical (due to the irregular shape of the boutons and stereological biases), to localise the estimated centres we first used spatial filtering (~100 nm range) and registered the area which has top 5% brightness in the hotspot: in the case of symmetrical hotspots this procedure generates exactly the same outcome as the standard Gaussian PSF-based localisation.

## ACKNOWLEDGEMENTS

This work was supported by the Wellcome Trust Principal Fellowship, European Research Council Advanced Grant, Medical research Council, Biology and Biotechnology Research Council (all UK), EXTRABRAIN Marie Curie Action (European Commission). The authors thank Mayeul Collot, University of Strasbourg, for generous donation of the Calcium Ruby-nano.

## AUTHORS CONTRIBUTIONS

T.P.J. designed and carried out imaging experiments and data analyses; K.Z. designed and implemented time-resolved imaging methods; N.C. carried out slice preparations, transfection and molecular biology protocols; J.S.M. and L.L.L. developed and provided SF-iGluSnFR glutamate sensor variants; T.P.J., K. Z. and D.A.R. analysed the data; D.A.R. narrated the study, carried out data analyses, and wrote the draft; all authors contributed to manuscript writing.

