## Supplementary Figures for "Multiplex imaging of quantal glutamate release and presynaptic Ca^2+^ at multiple synapses *in situ*"

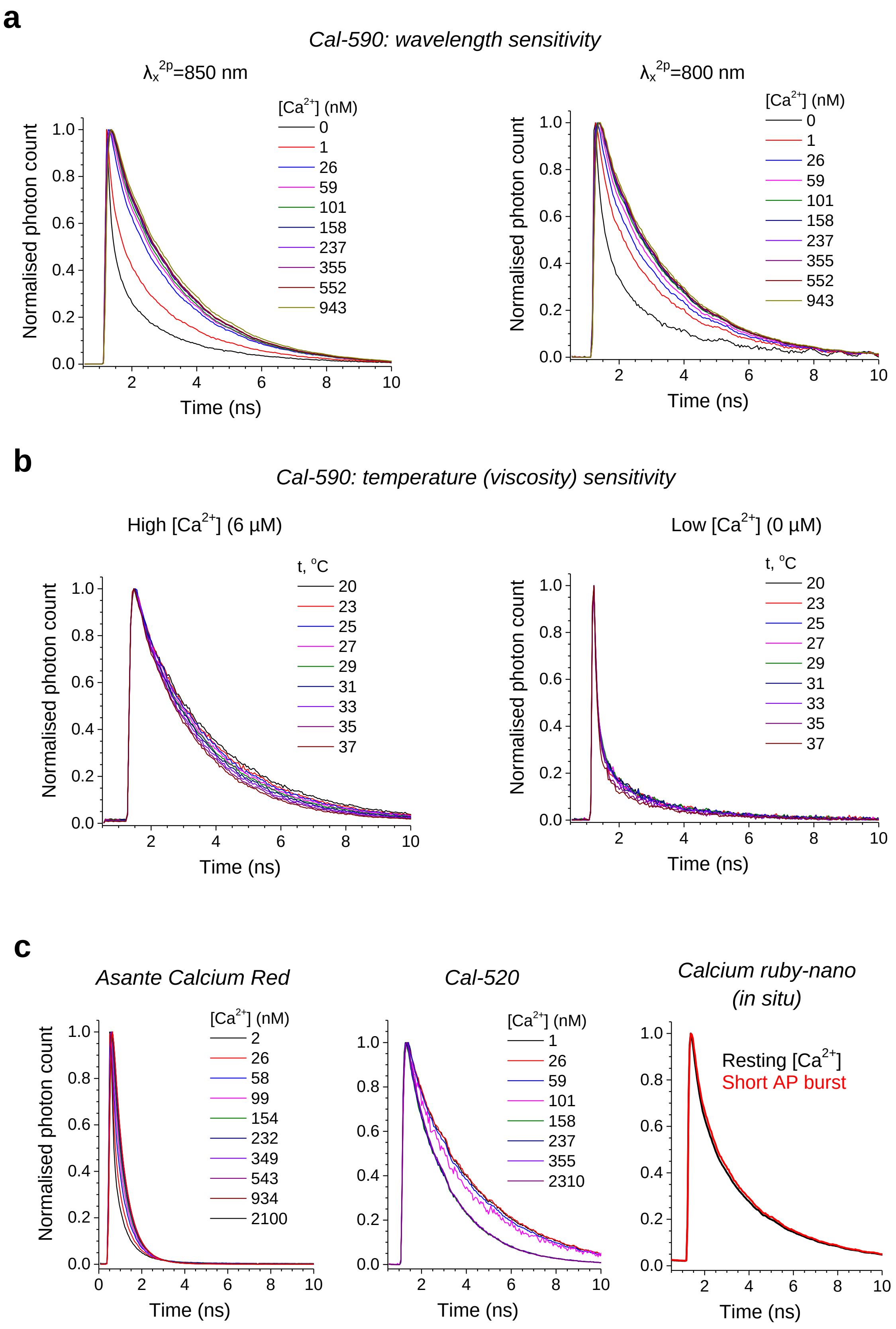

**Supplementary Figure 1. Cal-590 fluorescence lifetime sensitivity to excitation wavelength and temperature / viscosity.**

**(a)** Cal-590 lifetime sensitivity to $[Ca^{2+}]$ under different two-photon excitation wavelength, 850 nm and 800 nm, as indicated (temperature 33ºC). The detected sensitivity range for 0-200 nM $[Ca^{2+}]$ is narrower than that for the optimal excitation wavelength of 910 nm (Fig. 1a-b).

**(b)** Fluorescence decay of Cal-590 under saturating (left) and zero-clamped (right) $[Ca^{2+}]$, over the range of experimentally relevant temperatures, as indicated (viscosity).

**(c)** Testing the fluorescence decay $[Ca^{2+}]$ sensitivity for Asante Calcium Red, Cal-520, and ruby-nano, as indicated: the first two were evaluated using the standard FLIM calibration procedure as in **a-b** whereas Calcium ruby-nano was tested *in situ* (axonal bouton, CA3 pyramidal cell, organotypic hippocampal slices), by comparing its fluorescence decay in resting conditions (black) and during peak intensity response to a burst of four action potentials (at 20 Hz, red), as indicated; recordings at $\lambda_x^{2p}$ = 910 nm, 33ºC.

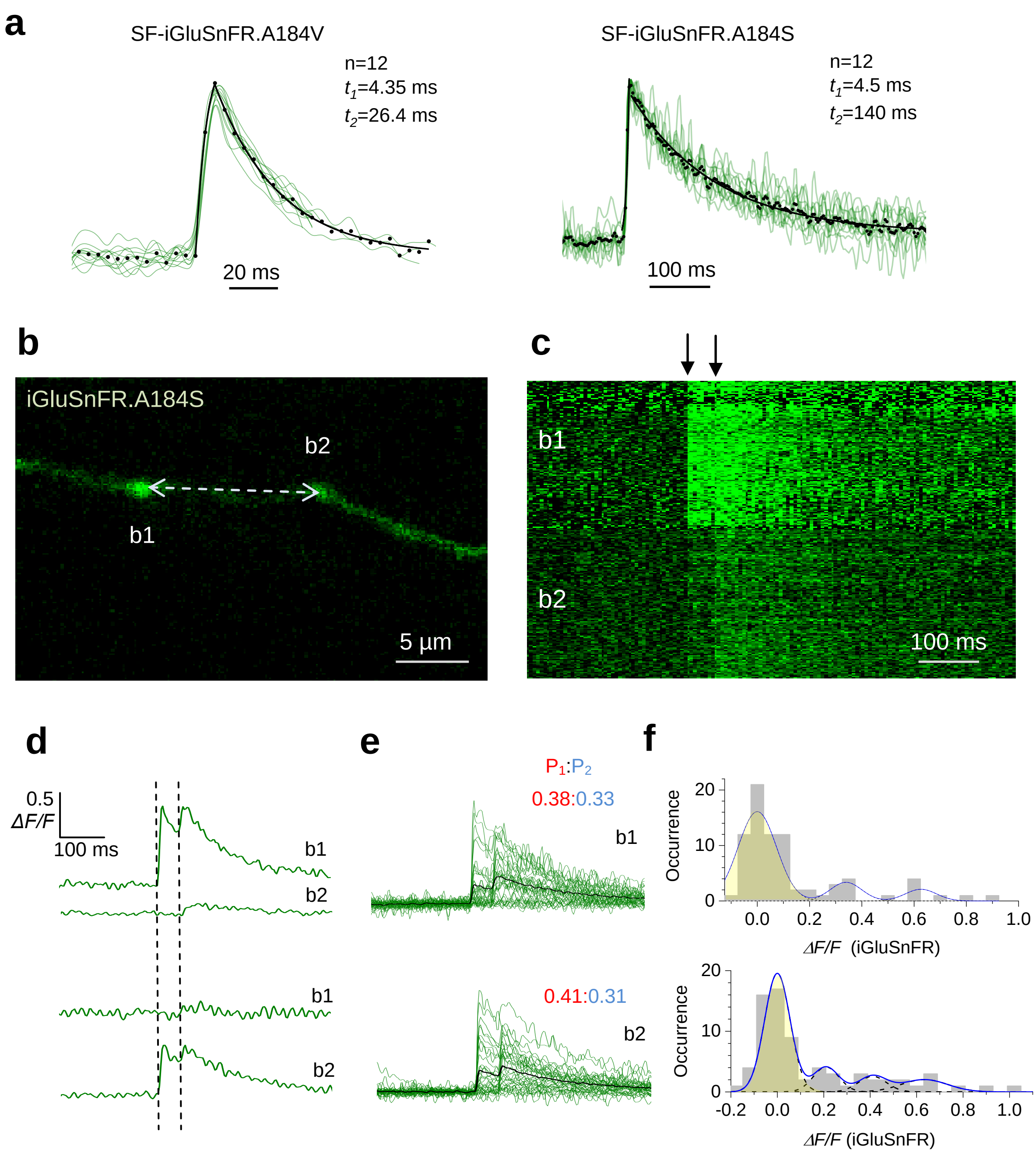


**Supplementary Figure 2. Simultaneous multi-synapse imaging of quantal glutamate release with iGluSnFR.A184S.**

**(a)** Comparative assessment of the binding (fluorescence) kinetics for SF-iGluSnFR.A184V and SF-iGluSnFR.A184S variants, as indicated, upon quasi-instantaneous glutamate release; green traces, normalised single-AP-evoked fluorescence responses recorded in individual boutons of CA3 pyramidal cell axons traced to area CA1; dots, all-trace average; black line, best-fit double-exponent approximation: $t_1$, rise constant; $t_2$, decay constant; n, number of trials.

**(b)** CA3 pyramidal cell axon fragment in area CA1 showing two presynaptic boutons (b1-b2); two-headed arrow illustrates laser scan trajectory, with the scanning dwell points in the bouton centres.

**(c)** A pseudo-linescan image of SF-iGluSnFR.A184S signals recorded simultaneously (one sweep example) at two boutons shown in **a** as indicated, during somatic generation of two APs 50 ms apart (arrows); glutamate releases and failures can be seen; further detail in Fig 2b.

**(d)** Example of two consecutive recordings (top and bottom; *ΔF/F* SF-iGluSnFR.A184S) from two boutons, as indicated, to illustrate no fluorescence signal cross-talk between the boutons. P1:P2, average probability of the first (red) and second (blue) release events.

**(e)** A summary of 36 trials (1 min interval) in the experiment shown in **a-b**; green traces, single-sweep SF-iGluSnFR.184S *ΔF/F* intensity readout at the two bouton centres; black traces, all-sweep-average.

**(f)** Amplitude histograms (SF-iGluSnFR.184S *ΔF/F* signal, first and second response counts combined; pre-pulse 8 ms baseline subtracted), with a semi-unconstrained multi-Gaussian fit (blue line, Methods) indicating peaks that correspond to estimated quantal amplitudes; the leftmost peak corresponds to zero-signal (failure; yellow shade); dotted lines, individual Gaussians; arrows, average amplitudes (including failures) of the first (red) and second (blue) glutamate response.

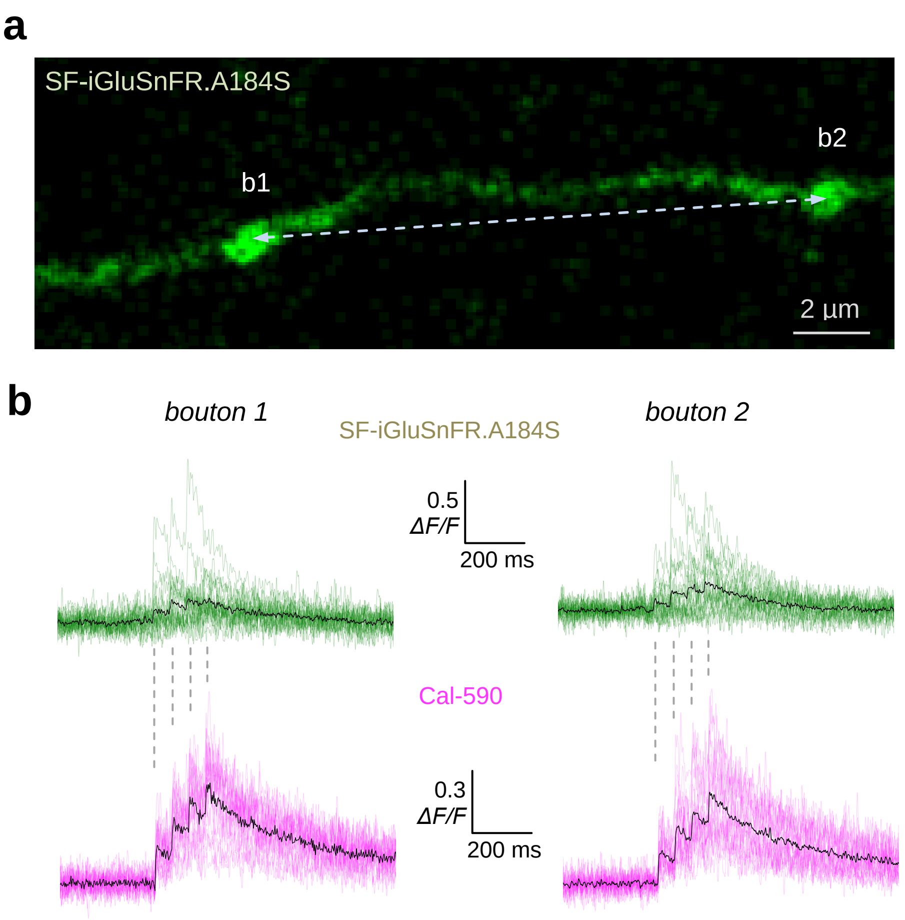


**Supplementary Figure 3. Simultaneous multi-synapse imaging of quantal glutamate release and presynaptic $Ca^{2+}$ dynamics with SF-iGluSnFR.A184S and Cal-590.**

**(a)** CA3 pyramidal cell axon fragment in area CA1 showing two presynaptic boutons (b1-b2; green channel); two-headed arrow illustrates laser scan trajectory, with the scanning dwell points in the bouton centres.

**(b)** A summary of 24 trials (1 min interval) in the experiment shown in **a**; green and magenta traces, single-sweep SF-iGluSnFR.A184S and Cal-590 *ΔF/F* intensity readout at the two bouton centres; black traces, all-sweep-average.

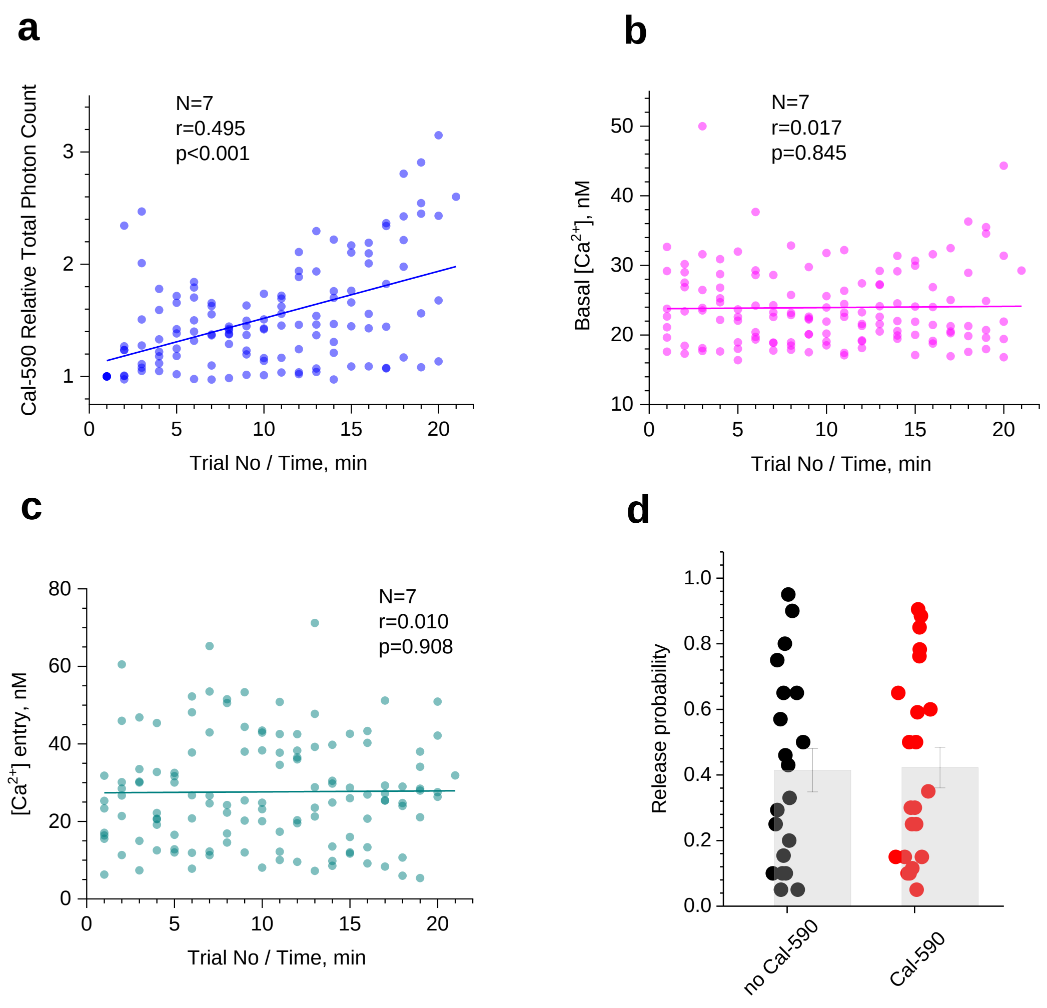


**Supplementary Figure 4. Intra-axonal $Ca^{2+}$ buffering by Cal-590 has little effect on [$Ca^{2+}$] FLIM readout or average release probability.**

**(a)** Total photon count for Cal-590 (fluorescence intensity, measured within 300 ms before the evoked four-AP train at 20 Hz, as in Figs. 3 and 4), normalised to the initial value, as recorded in individual axonal boutons (N = 7), plotted against the trial number (time). In these experiments, the intra-axonal Cal-590 concentration, hence $Ca^{2+}$ buffering capacity, continues to rise with time, before eventual equilibration; solid line, linear regression (r, Pearson correlation; p<0.001, slope significance).

**(b)** Presynaptic resting [$Ca^{2+}$] measured within 300 ms before the evoked four-AP train using Cal-590 FLIM readout, at axonal boutons shown in **a**, plotted against the trial number (time); other notation as in **a** (no correlation).

**(c)** An increment in presynaptic [$Ca^{2+}$] during evoked four-AP train measured with Cal-590 FLIM, at axonal boutons shown in **a-b**, plotted against the trial number (time); other notation as in **a-b** (no correlation).

**(d)** Probability of evoked glutamate release (in response to a single action potential) in axonal boutons of CA3 pyramidal cells (organotypic hippocampal slices) expressing iGluSnFr, with and without Cal-590 (300 μM) being loaded and equilibrated in whole-cell mode, as indicated. Dots, individual bouton recordings; bar graph, mean ± s.e.m. (0.41 ± 0.07 and 0.42 ± 0.06, n = 20 and n = 22, without and with Cal-590, respectively).

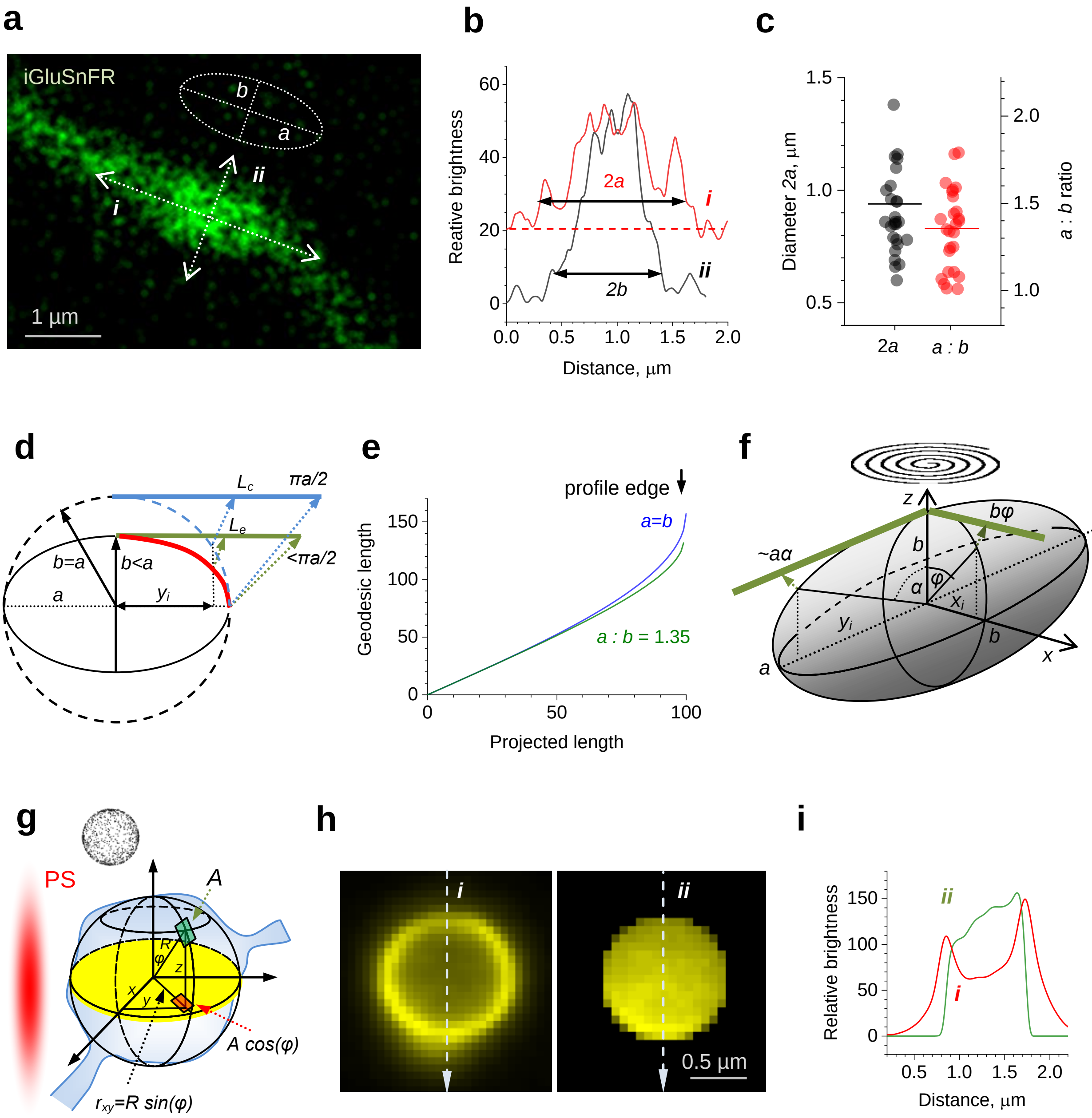


**Supplementary Figure 5. Stereological geodesic corrections for the signal spread and density on spherical or elliptical surfaces (of axonal boutons) projected onto the focal plane.**

**(a)** An axonal bouton (example) projection in the focal plane, with two profile sampling lines (*i* and *ii*) to estimate outline approximation by an ellipse (dotted oval), with *a* and *b* being major and minor axes, respectively.

**(b)** Brightness profiles along lines *i* and *ii* shown in **a**, as indicated. Two-headed arrows, effective width of the bouton along *i* and *ii* axes, to provide *a* and *b* values for the approximating ellipse, as shown. The cut-off values for *a* and *b* (arrow end positions) correspond to ~2SD of the background noise; horizontal dotted line, background reflecting the adjacent axon.

**(c)** Data scatters showing *a* values (mean ± SEM, 0.59 ± 0.02) and *a* : *b* ratios (1.35 ± 0.04) in the recorded sample of boutons (n = 26).

**(d)** Trigonometry diagram explaining geodesic corrections for distances on curved surfaces that are projected onto the (focal) *x-y* plane (as in Fig. 5d). For an elliptical section with major and minor axes *a* and *b*, respectively, the projected distance $x_i$ from the centre corresponds to the geodesic distance ($L_e$, green segment) which is consistently larger than that for a circular correction ($L_c$, blue segment).

**(e)** Comparison between $L_e$ and $L_c$ geodesic corrections (relative 1-100 scale), with the average *a* : *b* experimental ratio of 1.35 (as in **c**). The discrepancy between the two is small throughout the length reaching a maximum of ~15% towards the projection edge (elongated tip of the ellipsoid in **d**).

**(f)** Summary schematic for geodesic correction applied to a rotational ellipsoid (approximating the axonal bouton shape) projected onto a plane. For the running coordinate $x_i$, the corrected geodesic distance (along the *x* axis) is $b\varphi = b$ arcsin ($x_i / b$) whereas for the coordinate $y_i$, the geodesic distance (along the *y* axis) is ~ $a\alpha = a$ arcsin ($y_i / a$). Because the tornado scan (shown) is normally a circle inscribed into the bouton oval (Fig. 5e), there will be no data collected towards the tip of the ellipse, in the *y* direction (Fig. 5f). Thus, the ~15% discrepancy between circular and elliptic corrections which is seen towards the profile edge in the *y* direction (seen in **e**) will be effectively void, ensuring good approximation (estimated error <5%).

**(g)** Diagram illustrating stereological correction for the surface signal density on a spherical surface; inset, cartoon showing that the planar projection of spherical objects, with uniform surface distribution of fluorescence signals (dots), yields overestimated signal density towards the projection edges; red shade, approximate representation of the microscope's point-spread function (PSF). Sphere-like bouton structure is shown projecting onto the microscope focal plane (yellow)**.** In spherical coordinates (indicated), the differential (infinitesimal) element *A* of the sphere surface (green quadrangle) has an area of $r^2 \cdot sin(\varphi) d\varphi d\theta$ whereas its projection onto the $z$=0 plane (red quadrangle) has an area of $r^2 \cdot sin(\varphi) \cdot cos(\varphi)\ d\varphi d\theta$. The signal intensity in the projection is therefore boosted by factor $1 / cos(\varphi)$ compared to the surface intensity. Thus, applying the factor $cos(\varphi) = (1 - sin^2(\varphi))^{1/2} = (1 - r_{xy}^2 / r^2)^{1/2}$ to the image intensity provides the projection-corrected value.

**(h)** *Left,* focal plane projection of a nanoengineered microcapsule containing a fluorescent dye in its shell only (left) [1]. *Right*, the capsule shell fluorescence map corrected for the stereological bias as explained in **g**; dotted lines (*i, ii*), brightness profile sampling.

**(i)** Brightness profiles sampled in images shown in **h**, as indicated.
